# LARP6C regulates selective mRNA translation to promote pollen tube guidance in *Arabidopsis thaliana*

**DOI:** 10.1101/2020.11.27.401307

**Authors:** Elodie Billey, Said Hafidh, Isabel Cruz-Gallardo, Celso G. Litholdo, Viviane Jean, Marie-Christine Carpentier, Claire Picart, Katarina Kulichova, David Honys, Maria R. Conte, Jean-Marc Deragon, Cécile Bousquet-Antonelli

## Abstract

In angiosperms, non-motile sperm cells are delivered to the ovules for fertilization via a guided growth of the pollen tube. RNA binding proteins are key regulators that control mRNA fate post-transcriptionally and thus essential for normal cell function. But very little is known on the mechanistic bases of mRNA regulations and *trans*-acting factors governing male-female signalling. Here we demonstrate that the evolutionarily conserved RNA binding protein LARP6C is necessary for pollen tube guidance and fertilization. *larp6c* loss-of-function mutants exhibit male induced fertility defects as mutant pollen tubes frequently are unable to find ovules for fertilization. In mature pollen, LARP6C localises with a pollen specific poly(A) binding protein to cytoplasmic foci likely containing mRNPs. With RNA immunoprecipitation and sequencing, we demonstrate that LARP6C is associated *in vivo* with mRNAs required for pollen tube guidance or polarized cell growth. We further demonstrate using *in vitro* and *in planta* transient assays that LARP6C binds 5’-UTR box motifs to orchestrate the balance between translation, decay and storage of its mRNA targets. We propose a model where LARP6C maintains its mRNA target in translationally silent state likely to promote localized translation and guided pollen tube growth upon paracrine signalling.

## INTRODUCTION

The La Motif (LaM) is a structured RNA binding domain shared by hundreds of eukaryote proteins called La and Related Proteins (LARPs) which classify into five distinct subfamilies, amongst which the LARP6 group (Bousquet-Antonelli and Deragon 2009). In most LARPs, an RNA Recognition Motif (RRM1) is found immediately after the LaM, and forms a bipartite RNA binding unit called the La-module (Maraia et al. 2017). While the LaM is usually very well conserved between subfamilies, the RRM1s are specific to each subgroup (Bousquet-Antonelli and Deragon 2009; Maraia et al. 2017). Members of the LARP6 subfamily are found in Stramenopiles, Chlorophytes, plants, invertebrates and vertebrates (Bousquet-Antonelli and Deragon 2009; Merret et al. 2013b) and are characterized by the La-module and by a short conserved motif known as LSA at the C-terminus (<u>L</u>aM and <u>S</u>1 <u>A</u>ssociated) (Bousquet-Antonelli and Deragon 2009) that mediates protein-protein interactions (Weng et al. 2009; Vukmirovic et al. 2013; Cai et al. 2010b; Manojlovic et al. 2017). In mammals, LARP6 regulates the synthesis of type I collagen, an heterotrimer composed of two *α*1 and one *α*2 subunits whose assembly and correct secretion are dependent upon the coordinate translation of their transcripts at the endoplasmic reticulum (ER) (Zhang and Stefanovic 2016). The expression of type I collagen is primarily regulated by LARP6 binding through its La-module, to a stem loop in their 5’-UTRs (Cai et al. 2010a; Martino et al. 2015). LARP6 either stores or stabilizes the collagen mRNA transcripts in a translationally silent form by tethering them to vimentin filaments where they can further be activated for translation or subjected to decay (Challa and Stefanovic 2011). Alternatively LARP6 docks the mRNAs to non-muscle myosin filaments (Cai et al. 2010b; Manojlovic et al. 2017) and either recruits, via its LSA, STRAP (Serine Threonine kinase Receptor Associated Protein) (Vukmirovic et al. 2013) or the RNA helicase A (RHA) (Manojlovic et al. 2017), which respectively function as repressor and activator of translation. Through this mechanism, LARP6 orchestrates the coordinated translation of *α*1 and *α*2 collagen transcripts at the ER.

While LARP6 proteins are generally encoded by a single gene, vascular plant proteins are encoded by 3 to 6 genes and are of three evolutionary types, we named 6A, 6B and 6C (Merret et al. 2013b). Whilst members of the 6A subgroup most closely resemble LARP6 from other eukaryotes, B- and C-type orthologs carry, in addition to a reorganized La-module, a PABP-interacting motif 2 (PAM2). *Arabidopsis thaliana* has three *AtLARP6* genes, one from each type that we will refer to as LARP6A, B and C throughout this manuscript.

We previously demonstrated that LARP6C directly associates with the poly(A) binding protein (PABP) in plant, consistent with the presence of a PAM2 motif, and that its La-module binds to oligo(U_20_) homopolymers *in vitro*. In addition, in onion epidermis, LARP6C accumulates in the cytoplasm, the nucleoplasm and the nucleolus. Strikingly, LARP6C redistributes to stress granules (SGs) upon hypoxia associating herein with mRNP aggregates (Merret et al. 2013b). These data support the view that Arabidopsis LARP6C is an RNA binding protein likely to intervene in mRNA post-transcriptional regulations.

In flowering plants, the haploid male gamete (pollen grain) consists of a vegetative cell that encases two sperm cells, which are connected (through a so called cytoplasmic connexion) with the vegetative cell nucleus to form the male germ unit. A characteristic feature of angiosperm sexual reproduction is that sperm cells are non motile and must be carried simultaneously to the ovule by a pollen tube (Johnson and Preuss 2002). Upon adhesion to the stigma, the male vegetative cell hydrates, germinates and grows into a tube that delivers the two sperm cells to the ovule. The pollen tube is a greatly polarized and fast tip-growing cell which growth directionality is determined by various perceived cues. It is firstly guided in the gynoeceum by the pistil female sporophytic tissue (preovular guidance) then, upon exit from the transmitting tract, it navigates in gametophytic tissues guided by ovule-emitted signals (ovular guidance). Ovular cues guide the pollen tube alongside the funiculus to the micropyle. Upon reaching the ovule, the pollen tube stops growing and bursts to free the sperm cells that will fuse respectively with the egg cell (to form the embryo) and the central cell (to form the endosperm), hence completing the fecundation process (Higashiyama and Takeuchi 2015). In the pollen tube, upon reception and transduction of extracellular female signals, a complex cellular process targets the cell wall synthesis machinery at the site of perception, hence permitting the delivery of macromolecules at site of extension and the directional growth of the tube. This requires tip-localized receptors, flow of ions, intracellular trafficking of signalling molecules and proteins, but also vesicular trafficking, cytoskeleton-dependent transport and novel formation of cell wall through exocytosis (Feng et al. 2018; Hafidh et al. 2014; Higashiyama and Takeuchi 2015). The pollen tube growth also requires changes in the pattern of gene expression that at least in part rely on post-transcriptional mechanisms. Several data indeed suggest that all mRNAs required for germination and pollen tube growth are already present in the mature pollen grain and maintained in a translationally silent state until the progamic phase (Hafidh et al. 2018; Honys et al. 2000; Honys and Twell 2004; Scarpin et al. 2017). Nonetheless, how male gene expression is post-transcriptionally regulated to permit proper pollen guidance is largely unknown and no mRNA regulating factor required for proper pollen ovular guidance has been identified thus far.

In the present article we explore further the role of LARP6C and report that it is a pollen specific factor required for male fertility at the pollen tube guidance step. Our experimental results suggest that LARP6C orchestrates through direct binding the balance between translation, storage and degradation of transcripts that are required for pollen tube guidance. We propose that in dry pollen LARP6C could contribute to maintain its mRNA targets in a translationally silent state and permit their storage in mRNP granules. During pollen tube growth, LARP6C could act to stimulate the translation of at least some of its targets at the site of ovular cue perception

## RESULTS

### LARP6C is required for male fertility at the pollen tube guidance step

Western blotting and high-throughput transcriptomic analyses ((Klepikova et al. 2015; Schmid et al. 2005)) showed that, unlike LARP6A and 6B, LARP6C is almost exclusively expressed in pollen grains and pollen tubes (Figures 1A and S1). We therefore investigated its role in male fertility using two T-DNA insertion mutants, *larp6c-3* and *larp6c-4*. Western blotting showed that both alleles are loss-of-function (lof) mutants (Figure 1B). To establish whether *larp6c* is a gametophytic mutation, we monitored the transmission efficiency (TE) through the female (TE^f^) and the male (TE^m^) gametophytes by reciprocal crosses with wild type and heterozygous *larp6c-3/+* or *larp6c-4/+* mutants (Figure 1C). We found that both alleles were transmitted normally through the female gametophyte (TE^f^ *6c*-3 = 100% and *6c*-4 = 96.7%), whereas their transmission through the male was significantly reduced with *6c*-3 showing TE^m^ = 74.1% for *6c-3* and *6c-4* with TE^m^ = 68.2% (Figure 1C). This suggests that male but not female fertility is compromised by the loss of LARP6C function.

**Figure 1:**
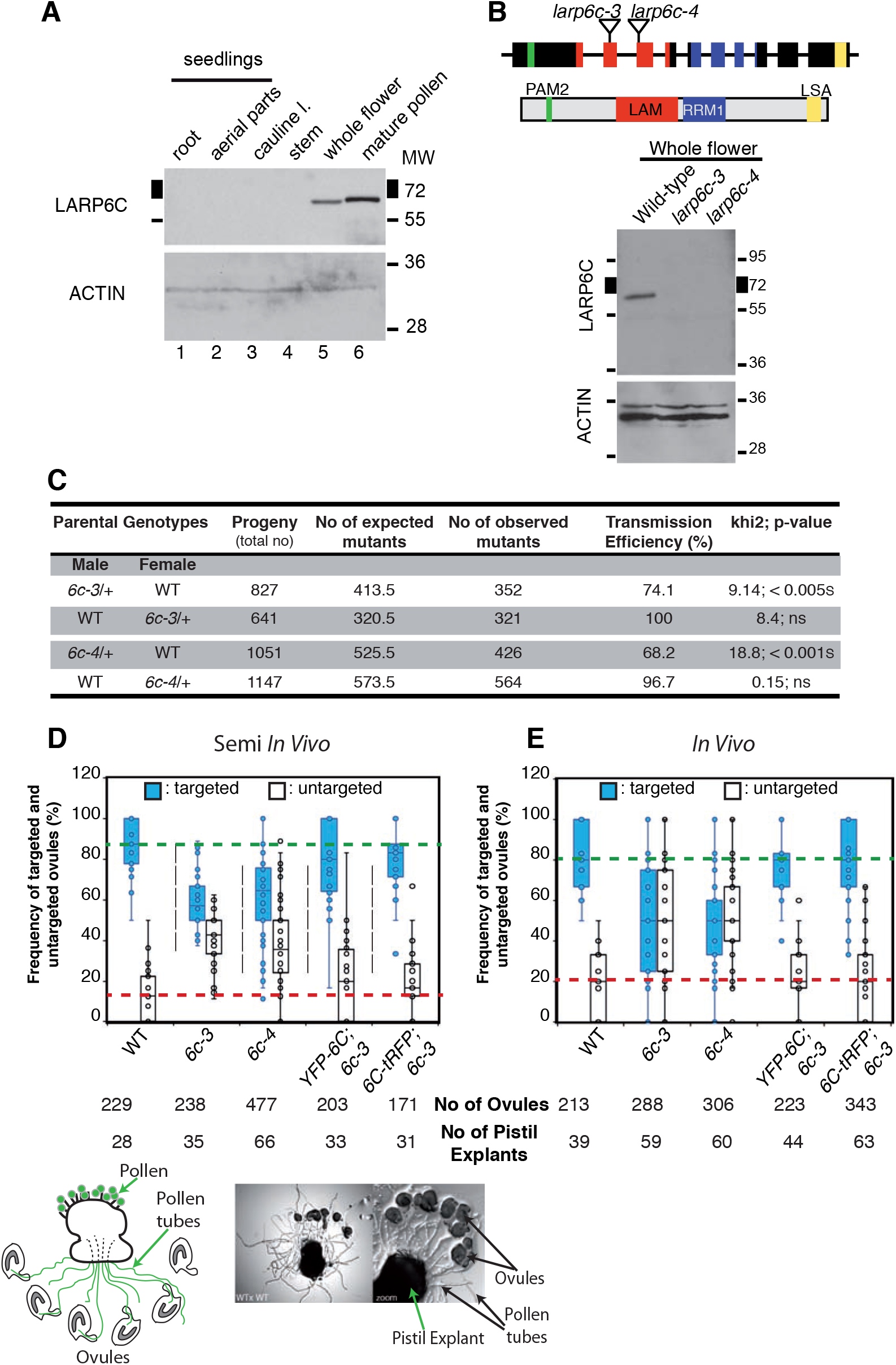
LARP6C is required for pollen tube guidance. **(A)**, Western blot analysis of steady-state LARP6C protein accumulation in seedling roots (lane 1) and aerial parts (lane 2), cauline leaves (lane 3), stem (lane 4), whole flower (lane 5) and mature pollen grain (lane 6). The same blot was probed with antibodies against LARP6C or ACTIN as loading control. **(B)**, *larp6c-3* and *larp6c-4* alleles are loss-of-function mutants. Top panel : schematic representation of the *LARP6C* gene and LARP6C protein. Plain boxes represent exons and lines represent introns. The color code for exons is identical to that for the protein conserved domains. Green: PAM2 motif, red: La Motif, blue: RRM1 and yellow: LSA. The insertion sites of the T-DNAs in *larp6c-3* and *6c-4* mutants are reported. Bottom panel: western blot analysis of LARP6C accumulation in flowers from wild-type, *larp6c-3* and *6c-4* lines. ACTIN is used as loading control. (**C**), Transmission efficiencies of *larp6c-3 and 6c-4* mutant alleles. Transmission Efficiency (TE) was calculated as: (([No of mutant]/[No of wild type]) x 100). ns: the number of mutant seedlings in the progeny is not significantly different from the expected number of mutant seedlings. (**D**), Semi *in vivo* (SIV) pollen tube guidance assays. Wild-type ovules were arranged around homozygous *ms*1 pistil explants pollinated with pollen from various homozygous backgrounds. The number of targeted (blue boxes) and non-targeted (white boxes) ovules was scored and results are represented as whisker boxplots. Above the graph are a cartoon and a representative pictograph of the setting up of a semi *In vivo* assay. (**E**), *In vivo* pollen tube guidance assays. Homozygous *ms*1 pistils were pollinated *in planta* with pollen from various homozygous genotypes, and the number of pollen tube-targeted (blue boxes) and non-targeted ovules (white boxes) were scored following aniline blue staining. Results are reported as whisker boxplots. On (D) and (E) the dotted lines respectively show the median values of targeted (green) and non-targeted (red) ovules by wild-type pollens.

There are different stages at which the male gametophyte could be deficient, including pollen development, germination, pollen tube growth or guidance and sperm delivery (Johnson and Preuss 2002). We monitored these pollen stages in *larp6c-3, 6c-4* and wild-type plants (Figures 1D, E and S2A-D). Assessment of male gametophyte maturation by DAPI staining revealed that 10% of the *6c-3* and *6c-4* mutant pollen grains did not complete their maturation and were arrested at the bicellular stage, compared to less than 3% in wild type (Figure S2A).

We next monitored the pollen germination and pollen tube growth using semi *in vivo* (SIV) ((Palanivelu et al. 2006) and experimental procedure) (Figure S2B, C) and *in vitro* assays (Figure S2D). We found that both *larp6c-3* and *6c-4* pollens behave as wild type for these two criteria, demonstrating that LARP6C is not required for pollen germination or pollen tube growth.

We next investigated LARP6C requirement for pollen tube guidance and ovule targeting competence (Johnson and Preuss 2002). We set up SIV experiments using pistils from *ms1* plants that are male-sterile mutant with normal function pistil (Van der Veen and Wirtz 1968) to assess the ability of *larp6c*-deficient pollen tubes to guide through the pistil tissues towards wild-type ovules for fertilization (Figure 1D). Whereas wild-type pollens target 80% to 100% of the ovules (with a median value of around 90%), both *larp6c-3* and *6c-4* pollens showed a significant reduction, targeting 50 to 70% of the wild type ovules (with a median value of 60%) (Figure 1D). This failed pollen tube guidance phenotype was complemented by the N- and C-terminal LARP6C fusion constructs (YFP-LARP6C and LARP6C-tRFP) (Figure 1D and Figure S3A-B). In the competition assays irrespective of the order of pollination or the side of the stigma in which the pollen grains were dusted on, the *larp6c-3* and 6c-*4* pollens also showed a clear deficiency in ovule targeting with a maximum targeting efficiency of 20% from the expected 50% rate (Figure S2E). This further supports that the observed guidance phenotype directly relates to *larp6c* deficiency. We further monitored the ability of the *larp6c* and *larp6c*;LARP6C-tRFP complemented pollen to target ovules *in planta* (Figure 1E). We hand pollinated *ms1* pistils and following aniline blue staining, scored the number of targeted and non-targeted ovules. Here again the *larp6c* deficient pollen tubes displayed only 60% ovule targeting efficiency versus the 80% average observed in wild-type pollinated pistils. The targeting efficiency by the *larp6c*;LARP6C-tRFP complemented lines recovered to a wild-type frequency (Figure 1E).

To verify that *larp6c* lof specifically affects male fertility, we examined the capability of *larp6c-3* and *6c-4* mutant ovules to attract wild-type pollen tubes by semi *in vivo* and *in vivo* assays and found no significant difference from wild type (Figure S4). In summary, our results support that LARP6C is required for male fertility at least in part through its function during pollen tube guidance.

### LARP6C forms dynamic cytoplasmic granules in mature pollen grains

To get a better understanding of LARP6C cellular roles, we firstly monitored its subcellular distribution across pollen maturation and during pollen tube growth (Figure 2 and S3D). In microspores (UNM), LARP6C is only detected in the nucleolus. At the bicellular stage (BCP) LARP6C is much more generally distributed, as it can be detected in the cytoplasm of the vegetative cell (VC) as well as in the vegetative and generative cell nuclei (VCN and GCN). At the tricellular stage (TCP) LARP6C subcellular distribution is similar to that at the bicellular stage, with possibly reduced amounts of LARP6C in the sperm cell nucleus (SCN) compared to other subcellular compartments. In mature pollen grain (MPG) LARP6C displays a complex localization pattern. It is clearly present in the VC cytoplasm, but transitioned from a diffuse pattern to a granular, aggregate-like pattern. In addition to this granular distribution, LARP6C appears to concentrate around the membrane surrounding the vegetative cell nucleus, the sperm cells (SCs) and the cytoplasmic connection (CC) linking the SC1 to the VCN. In pollen tubes, LARP6C no longer showed the granular pattern and is clearly excluded from the SCs.

**Figure 2:**
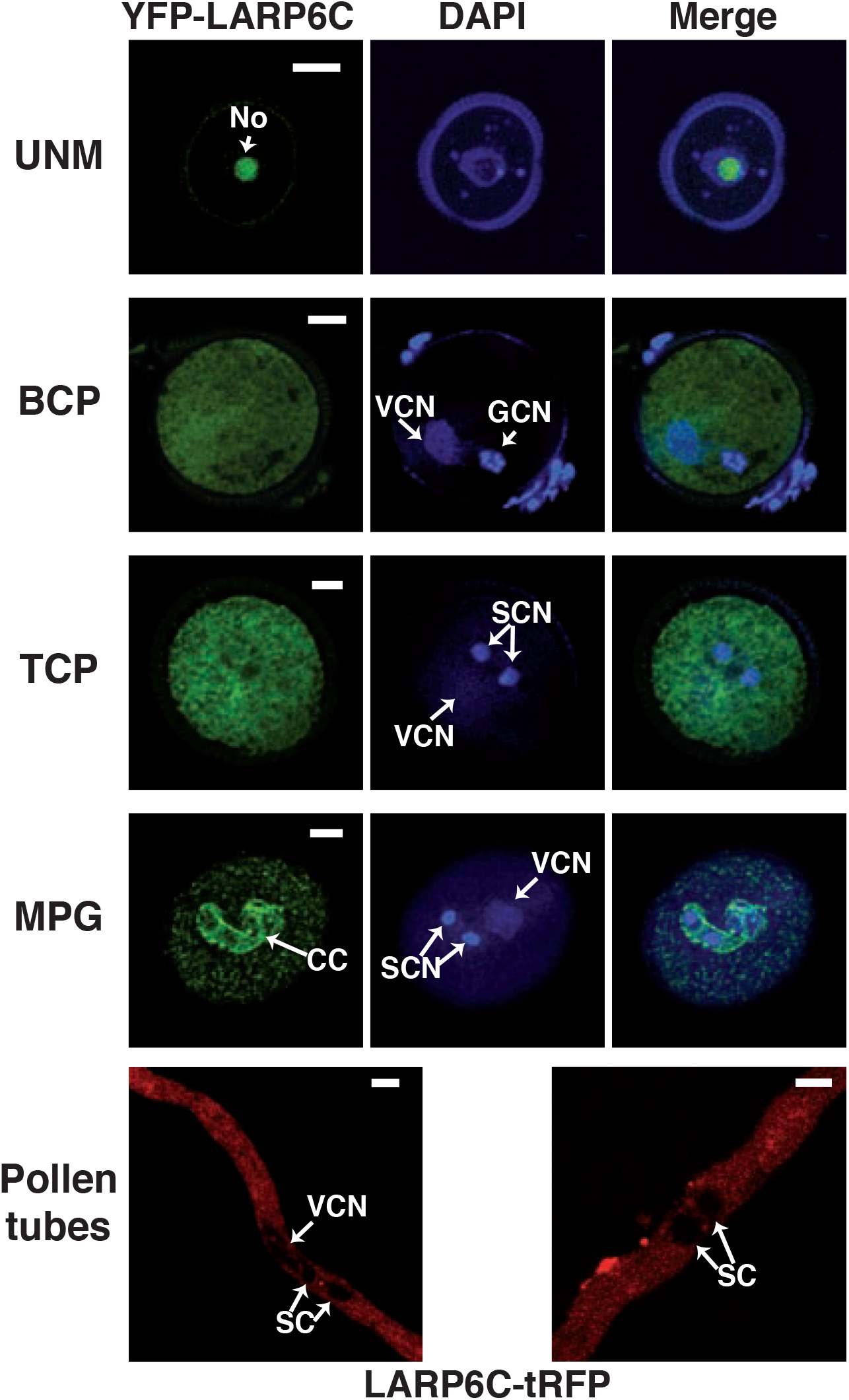
LARP6C shows a dynamic subcellular distribution in pollen. Subcellular localization of fluorescently tagged YFP-LARP6C by confocal microscopy at different stages of pollen maturation: UNM : uninucleate microspore, BCP: bicellular pollen, TCP: tricellular pollen, MPG: mature pollen grain and Pollen tube. Arrows indicate the position of the Nucleolus (No), Generative Cell Nucleus (GCN), Vegetative Nucleus (VCN), Sperm Cell Nucleus (SCN) and Cytoplasmic Connection (CC). Scale bars correspond to 5 μM. Representative images of LARP6C fused at its C-terminus with a tagRFP reporter are shown in supplemental Figure 2D.

To determine more precisely the distribution of LARP6C in mature pollen grains, colocalization experiments between LARP6C-tRFP and various markers were undertaken (Figure 3). The histone H2B-GFP fusion protein, expressed under the control of the tomato pollen-specific *LAT52* promoter (Twell et al. 1989), was used to label the VCN, thereby demonstrating the absence of LARP6C in this compartment. GEX2 is a sperm cell-expressed transmembrane protein (Mori et al. 2014) that labels the SC membranes and the cytoplasmic connection between SC1 and the VCN (Brownfield et al. 2009; McCue et al. 2011; Mori et al. 2014). Confocal analyses showed that LARP6C and GEX2 display distinct distributions. The tRFP signal is surrounding the GFP-GEX2 signal decorating the membrane of the male germ unit but not the membrane connecting the two sperm cells, supporting that LARP6C is excluded from the sperm cells and therefore only resides in the VC cytoplasm in mature pollen.

**Figure 3:**
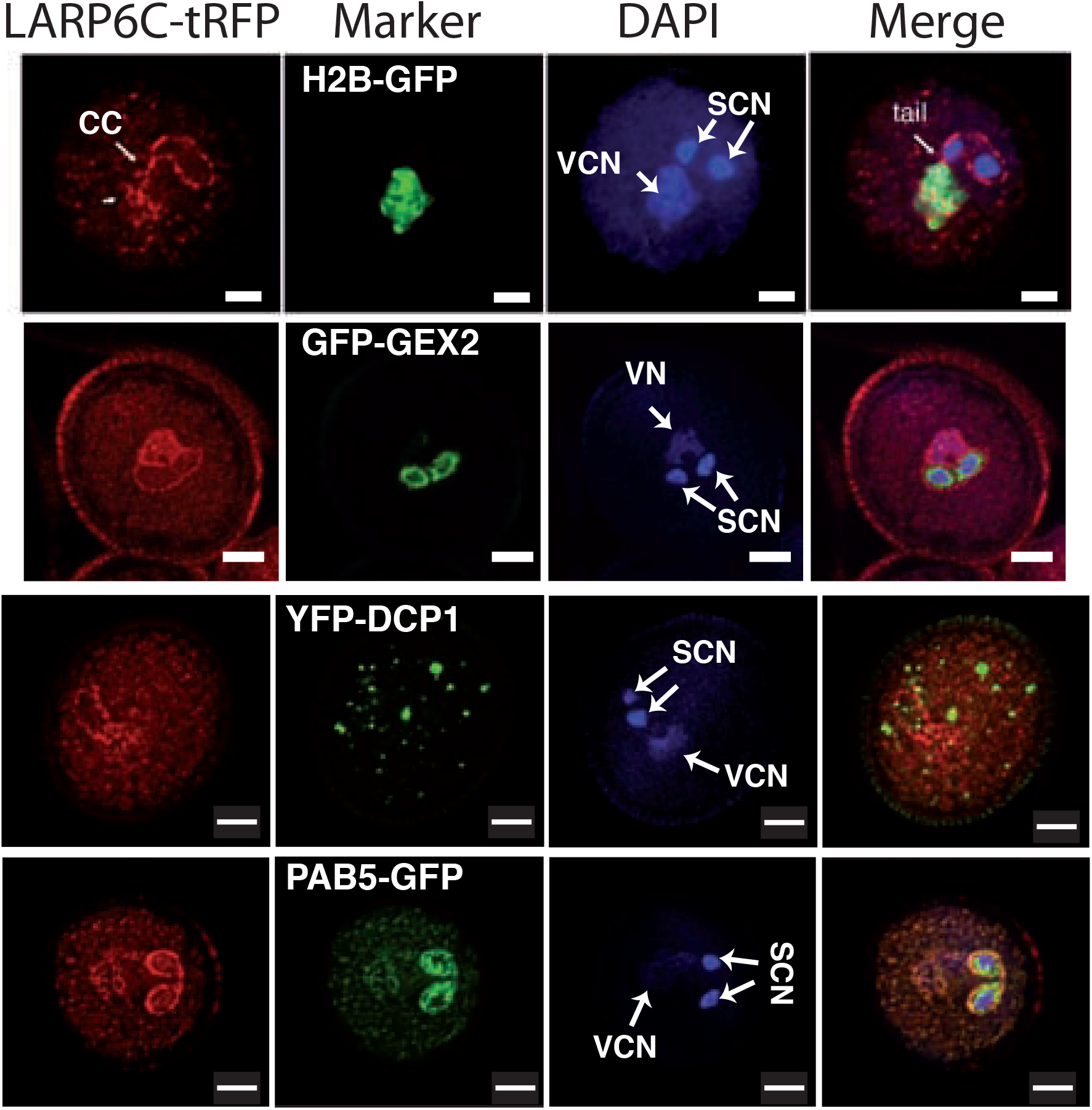
LARP6C forms mRNP aggregates that contain the Poly(A) Binding Protein in the vegetative cell cytoplasm. Confocal observation of mature pollen grains that stably coexpress the LARP6C-tRFP fusions and: the H2B-GFP fusion, the GFP-GEX2 fusion, the YFP-DCP1 fusion or the PAB5-GFP fusion. Except for the H2B-GFP marker which is expressed from the tomato *LAT52* promoter, all marker genes are expressed from their own upstream genomic sequences. Arrows indicate the position of Vegetative Cell Nucleus (VCN), Sperm Cell Nucleus (SCN) and Cytoplasmic Connection (CC). Scale bars correspond to 5 μM.

We next inquired on the nature of the LARP6C cytoplasmic foci (Figure 3). Eukaryotic cells contain many types of membrane-less cytoplasmic aggregates formed of translationally silent mRNAs and RNA binding proteins (RBPs) (Buchan 2014). So far only two types of mRNP granules have been identified in plant cells: processing bodies (p-bodies) and stress granules (SGs) (Weber et al. 2008). Plant p-bodies are - as in other eukaryotes - characterized by the presence of actors of the general mRNA turnover process, such as the DCP1/DCP2 decapping holoenzyme, the XRN4 exoribonuclease or the deadenylation complex. Stress granules on the contrary contain factors belonging to the translational machinery, amongst which the poly(A) binding protein (Weber et al. 2008). First clues on the nature of LARP6C aggregates observed in mature pollen were derived by colocalization experiments with specific markers of p-bodies and SGs respectively, namely DCP1 and PABP. We used a YFP-tagged DCP1 protein expressed from its own upstream genomic sequences (Merret et al. 2013a). Although YFP-DCP1 displays a punctuate distribution in the VC cytoplasm, similarly to LARP6C, no obvious colocalization of DCP1 and LARP6C was observed. Of note, DCP1 accumulation appears to be restricted to the VC cytoplasm, at least in mature pollen and contrary to LARP6C, it does not localize at the periphery of the sperm cells or the vegetative nucleus. The Arabidopsis genome contains seven genes (*PAB2* to *8*) coding for canonical PABPs, including *PAB3, 5, 6* and *7* whose transcripts accumulation is restricted to pollen (Honys and Twell 2004; Belostotsky and Meagher 1996; Belostotsky 2003; Chekanova et al. 2001; Chekanova and Belostotsky 2003). We fused the genomic sequence of the *PAB5* gene (the most expressed of all four pollen specific *PABP* genes) to a YFP tag under the control of its own upstream sequences. In the VC cytoplasm, the YFP-PAB5 protein, displayed a pattern identical to that of the LARP6C protein; it accumulated in foci, surrounding the sperm cells and was excluded from the vegetative nucleus, but, contrary to LARP6C, it is also present in the cytoplasm of the sperm cells.

In summary, our data indicate that LARP6C shows a dynamic subcellular distribution across pollen maturation. We found that in mature pollen LARP6C is restricted to the vegetative cell cytoplasm and is present in aggregates that contain at least one actor of the translation process.

### RIP-seq pulldown revealed LARP6C mRNA targets in mature pollen grain

To explore LARP6C molecular functions we sought to identify its mRNA targets using an unbiased RNA immunoprecipitation and sequencing (RIP-seq) strategy. We conducted the RNA immunoprecipitation procedure from complemented *larp6c-3* lof mature pollen grains expressing a proLARP6C-FlagHA tagged version (6C-FH here on) (Figure S3A,C) and from wild-type pollen grains as negative control. After assessing the efficiency of the immunoprecipitation through western blotting (Figure S5A), RNAs were extracted from the input and eluate fractions and subjected to RNA-seq. After mapping and filtering of the dataset (see supplementary data procedure) a rpkm (read per kilobase per million mapped reads) value was calculated for each protein coding gene. The replicates were found to be reproducible (Figure S5B), enabling us to build table with a mean normalized value per gene and filtered out those that do have at least 1 rpkm value in one of the four samples (input or eluate from wild type or 6C-FH) (Table S1). With such criteria we retained a list of 7356 expressed genes that is expected for a pollen transcriptome which is less complex than that of sporophytic tissues (Loraine et al. 2013).

To find putative LARP6C mRNA targets, we first filtered out transcripts with Input-WT/Input-6C-FH ratio >1.2 in order to select for transcripts whose levels did not vary between control and 6C-FH inputs. Then we picked transcripts with an RE (RE is the eluate over input ratio) value in the 6C-FH samples above 1, to select for mRNAs that were enriched by the immunoprecipitation procedure (see Figure S5C for a diagram of the filtering workflow). To further increase stringency and reveal mRNAs that are most likely to be direct targets of LARP6C, we compared the RE_WT_ to the RE_6C-FH_ and, based on the repartition of the values (Figure S5D), we selected transcripts with an (RE_6C-FH_/RE_WT_) value of 3 or more. Through above series of filters, we identified 115 genes as direct or indirect mRNA targets of LARP6C (Table S1).

No gene ontology (GO) term was found to be significantly enriched amongst the 115 mRNAs identified as potential LARP6C targets. Nonetheless a gene to gene analysis of the proteins they encode shows that 25% code for factors that are membrane associated, 25% relate to sensing and signalling and 25% to membrane synthesis (Table S1). In addition several genes are known to be involved in polarized tip growth. The MPK6 kinase (Guan et al. 2014) and the TFIIB (Zhou et al. 2013) general transcription factor are necessary for pollen tube guidance. Exo70H3 is a subunit of the exocyst secretory complex essential for plasma membrane nanodomains specification that define sites for pollen tube directional growth (Li et al. 2010; Synek et al. 2006; Vukasinovic and Zarsky 2016). WAV2 participates in the regulation of Arabidopsis root directional growth in response to environmental cues (Mochizuki et al. 2005) and the AT3G50910 protein is a homolog of human DCC (Delected in Colorectal Cancer), a transmembrane receptor of Netrin, an evolutionarily conserved chemical cue of developing axon guidance. Several other LARP6C RIP targets could be required in pollen tube guidance and growth by acting at the signal perception/transduction levels, including PIRL5 (Forsthoefel et al. 2005), CIPK1 and 14 (Mao et al. 2016) and AT3G54800 which bears a PH and a START lipid binding domains and could therefore be involved in lipid signalling (Tang et al. 2005). Finally, the list of LARP6C targets also includes mRNAs coding for factors involved in cell wall synthesis and organization. This comprises actors of sphingolipid metabolism affecting pollen tube guidance (Tartaglio et al. 2017), such as the DES1-like sphingolipid Δ4-desaturase (Michaelson et al. 2009) and LCB2 which is involved in the first step of sphingolipid biosynthesis (Teng et al. 2008). Players in tri- or diacylglycerol metabolism, the homeostasis of which is necessary for correct pollen tube growth (Botte et al. 2011; Pleskot et al. 2012) are also present; including TAG1 that catalyzes the first step of the triacylglycerol synthesis pathway, SDP1, a triacylglycerol lipase and MGD2 that catalyses the formation of monogalactosyldiacylglycerol (MGDG) from diacylglycerol (DGDG) (Botte et al. 2011). Strikingly, 66% of the RIP-identified LARP6C targets are enriched or specifically expressed in pollen as compared to their expression in seedlings, while only 33% of the whole pollen transcriptome shows such higher or specific expression (Loraine et al. 2013).

### LARP6C directly binds to the 5’-UTRs of its mRNA targets

RNA binding proteins often bind to primary sequence motifs shared between their mRNA targets to regulate their fate. Considering that *cis* regulatory elements are mostly located in the transcripts 5’ and/or 3’ untranslated regions (UTRs), we looked for primary sequence motif(s) shared between the UTRs of the putative LARP6C targets. Using the Araport database, we found and retrieved the 5’- and 3’-UTRs for 112 genes and ran a discriminative motif search using a MEME search (at the MEME suite portal (https://meme-suite.org/). Whereas no significant motif could be detected in the 3’-UTRs of LARP6C mRNA targets, we identified two regions we named A-box (E^value^: 5.5E^-99^) and B-box (E^value^: 9.3E^-91^) motifs that are highly enriched in the 5’-UTRs (Figure 4A). As a control we ran an identical motif search using a set of 112 randomly chosen 5’- and 3’-UTRs from genes expressed in pollen according to our transcriptomic analyses. We found no significant enrichment for an A- or a B-type region, supporting that these sequence motifs are specific for mRNAs that are in a complex with LARP6C.

**Figure 4:**
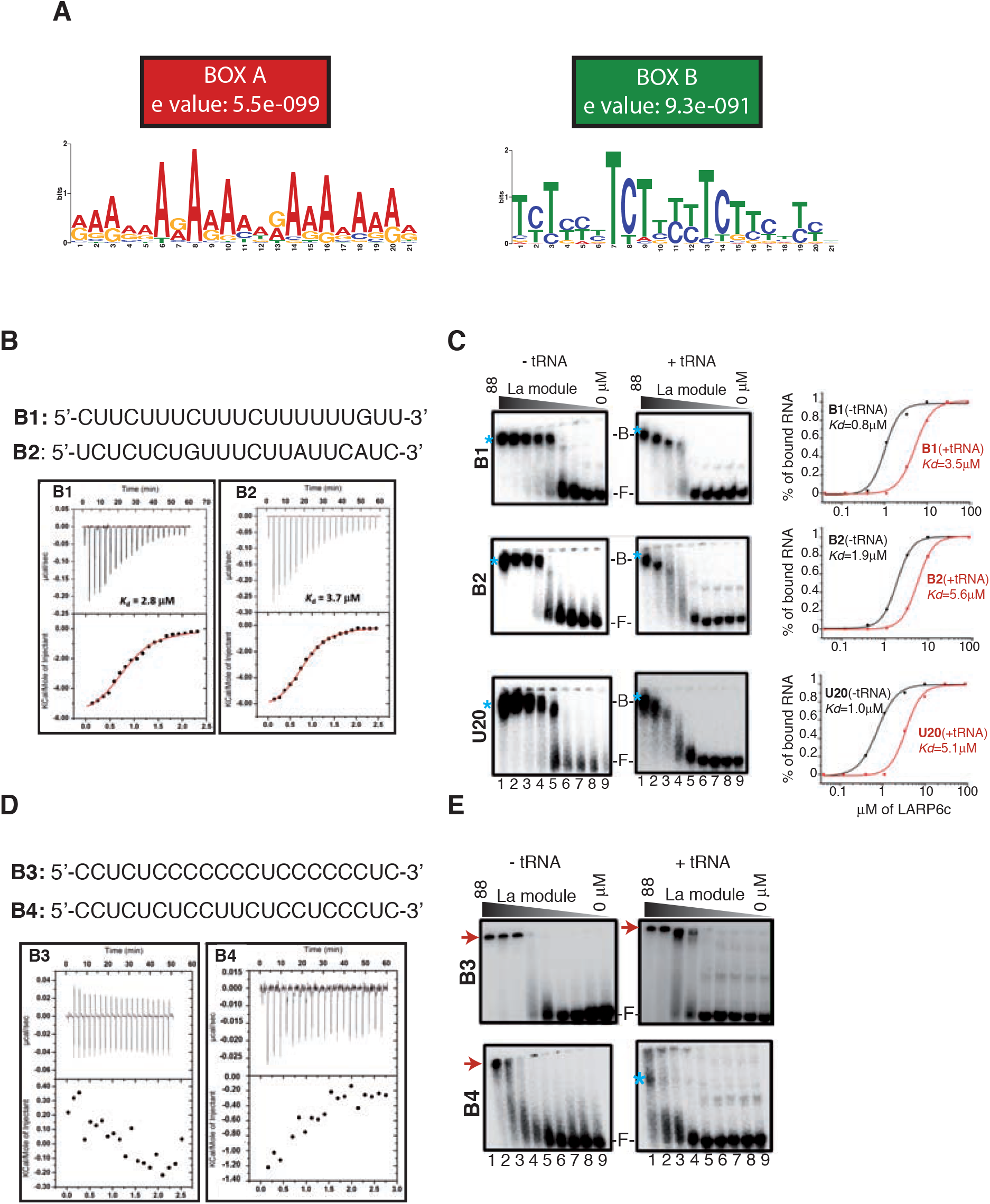
The LARP6C La-module binds the B-type RNA boxes. **(A)**, Consensus sequences of the conserved motifs and their approximate location (add the graph please shared by the 5’-UTRs of LARP6C mRNA targets. Calorimetric **(B, D)** and EMSA **(C, E)** analyses of the interaction between the LARP6C La-module (encompassing residues 137-332) and oligos B1, B2 **(B, C)**, U_20_ **(C)**, B3 and B4 **(D, E)**. In **(B)** and **(D)**: for each graph, the upper panel corresponds to the raw titration data showing the thermal effect of injecting an RNA oligo solution into a calorimetric cell containing the recombinant LARP6C La-module. The lower panels show the normalized heat for the titrations obtained by integrating the raw data and subtracting the heat of the RNA dilution. The red lines on graphs for oligos B1 and B2 **(B)** represent the best fit derived by a non-linear best-square procedure based on an independent binding site model. The dissociation constants (Kd) are indicated for the B1 and B2 oligos, the thermodynamic parameters are shown in Table S2. **(C, E**), EMSA analyses of LARP6C La-module binding to: B1, B2 or oligo U_20_ (C), B3 or B4 **(E)**. Decreasing concentrations (μM) (88 (lane 1), 29.3 (lane 2), 9.8 (lane 3), 3.3 (lane 4), 1.1 (lane 5), 0.4 (lane 6), 0.12 (lane 7), 0.04 (lane 8) and 0 (lane 9)) of the recombinant LARP6C La-module were mixed with 3nM of 5’-labelled oligos. B stands for Bound, F for Free, blue asterisks mark the RNA-protein complex and the red arrows on panels E shows samples retained into the gel wells. Experiments were conducted in absence (-tRNA) or presence (+tRNA) of unlabelled competitor (tRNAmix of *E*.*coli* MRE 600 at 0.01 mg/mL concentration). Graphs on panel C show the quantification of the bound RNA fraction versus the protein concentration in the absence (black lines) or presence (red lines) of tRNA competitor. The values of the dissociation constants are reported. Kd reported for the EMSA experiments were calculated out of three independent replicates with the following standard deviations: LARP6C-B1 (-tRNA): 0.8 +/- 0.1; LARP6C-B1 (+tRNA): 3.5 +/- 0.3; LARP6C-B2 (-tRNA): 1.9 +/- 0.3; LARP6C-B2 (+tRNA): 5.6 +/- 0.2.

The A motif is a purine-rich 21 nt sequence, with a clear dominance of A over G nucleotides while the B motif is a pyrimidine-rich 21 nt sequence with a majority of U (T in DNA) over C repeats (Figure 4A, Table S1). A closer examination of sequences that contain the A and/or B consensus revealed two major categories for each box type. In particular, sequences harboring the A motif are mainly composed of either A or AG/AAG repeats whilst, sequences sharing the B consensus motif contain either U or UC/UUC repeats (Table S1). Amongst the 112 identified 5’-UTRs, 70 (62.5%) bear the A motif, 74 (66%) the B motif and 49 (43.7%) carry both (Figure S5E).

The presence of motifs shared by transcripts co-immunoprecipitated with LARP6C *in vivo* suggests that one or both of these boxes may mediate direct binding of LARP6C. To challenge this hypothesis we tested the RNA binding unit of LARP6C, the La-module, for its ability to interact with these boxes *in vitro* (Figures 4B-E and S6). We designed two oligonucleotide sequences for the A motifs, called A1 and A2 representing the “A-rich” and the “AG/AAG-rich” types respectively (Figure S6A). Analogously B1 and B2 were designed for the B motifs exemplifying the “U-rich” and “UC/UUC-rich” types respectively (Figure 4B). As a control, we designed oligos B3 and B4 representing respectively C- or CU/CCU-rich sequences. A recombinant version of the LARP6C La-module, identical to the one used in Merret *et al*. (Merret et al. 2013b), was expressed as previously described in *E. coli* and its ability to bind the A and B type oligo RNAs was quantified by isothermal titration calorimetry (ITC) and electrophoretic mobility shift assay (EMSA) (Figure 4B-E and S6). In the ITC experiments serial titration of LARP6C La-module with RNA oligos B1 and B2 generated in both cases a profile that could be fitted to a sigmoid-shaped binding curve centered around a 1:1 stoichiometry, with a dissociation constant in the low micromolar range (Kds of 2.8 μM and 3.7 μM respectively), suggesting a high affinity of LARP6C La-module to B1 and B2 type motifs (Figure 4B and Table S2). The binding mode of LARP6C La-module for the B1 and B2 oligos is similar (but with higher affinity) to that we previously observed for an U20 homopolymer (Table S2 and (Merret et al. 2013b)). On the contrary, oligos B3, B4, A1 and A2 revealed no binding to the recombinant LARP6C domain by ITC in the same experimental conditions used (Figure 4D, Figure S6A and Table S2). Parallel EMSA assays were performed to confirm ITC results, using RNA oligos labelled at the 5’ with γ-^32^P, to monitor RNA-protein complex formation in native gel electrophoresis. Experiments were conducted in the presence or absence of tRNA competitor. For oligos showing the ability to form a complex with the La-module, the fraction of bound RNA was plotted against the protein concentration and the dissociation constant determined (Figure 4C). In agreement with ITC, EMSA experiments confirm that B1 and B2 oligos associate with LARP6C La-module, with estimated K_d_s in the low micromolar range (Kds of 0.8 μM and 1.9 μM respectively) suggesting here again a high affinity of LARP6C La-module towards B1 and B2 oligos (Figure 4C). B2 binding behaviour is very similar to what we observed for an oligo(U_20_) whereas B1 shows a slightly higher binding affinity. Assays with oligos A1, A2, B3, oligo(C_20_) and to a lesser extent with B4 were hindered by samples being retained in the wells of the gel at high protein concentration (Figure 4E and S6B), an issue most probably linked to protein aggregation (Hellman and Fried 2007). Despite changing various experimental conditions, this issue could not be overcome. The presence of a tRNA competitor marginally helped in the case of B4 albeit association with LARP6C was observable only at high protein concentration and a Kd could not be calculated as the binding curve did not reach a plateau (Figure 4E).

Collectively these experiments demonstrate that LARP6C La-module binds to oligos B1 and B2 with high affinity and with a similar association mechanism observed with oligo(U_20_). B1, which contains three stretches of U residues, one of which 6 uridine-long, appears to bind slightly tighter than B2 and U_20_. Altogether these results indicate that the LARP6C La-module strongly binds to B type motifs and does not associate with A type motifs, at least *in vitro*, and that it displays low affinity for B4.

We hence propose that *in vivo* LARP6C directly binds to mRNAs that carry in their 5’-UTRs, a B1 or B2-type box, to impose posttranscriptional regulation on its mRNA targets.

### LARP6C is not required for accumulation of its mRNA targets in mature pollen grain

To have a global view of LARP6C mode of regulation on its mRNA targets’ accumulation we monitored the polyadenylated transcriptome of mature pollen grain in the absence of LARP6C. Pollen grains were collected from wild-type and *larp6c-3* plants grown together in a greenhouse (see supplemental experimental procedure), total RNAs extracted and the polyadenylated fraction subjected to next generation sequencing. Following mapping and filtering of the reads (see supplementary experimental procedure), we retained a list of 7399 transcripts with at least 1 rpkm as reliably expressed from both Col-0 and *larp6c-3* (Figure S7, Table S3). We then used a DE-seq pipeline to find genes whose transcripts differentially accumulate in the mutant background and retained those with a fold change of at least 1.5 between wild type and *larp6c-3*. We found 2174 genes either up- or downregulated in *larp6c-3* pollen (Figure 5A). Monitoring the fold changes of these transcripts between wild type and *larp6c-3*;6C-FH in the input fraction of the RIP-seq samples, we observed that they are centered around 1 (Figure 5B). This suggests that the expression of the LARP6C-FlagHA transgene in the *larp6c-3* background fully complemented the *larp6c-3* mutation and restores a wild-type level expression, proving that the LARP6C-FlagHA translational fusion is functional and that the misregulation of *larp6c-3* pollen transcriptome is biologically significant and the consequence of the loss of LARP6C function. We then asked if the steady-state levels of LARP6C targets are affected in *larp6c-3* pollen grains. Of the 115 targets, 96 were detected in the RNA-seq data and only 15 of these differentially accumulated in the *larp6c-3* background (Figure 5C, Table S3). Of these ten are downregulated by 1.5 to 2.8 fold and five are upregulated by 1.5 to 1.7 fold in *larp6c-3* (Figure 5D). These observations suggest that, at least in dry pollen, LARP6C is not required for maintaining steady-state levels of its mRNA targets.

**Figure 5:**
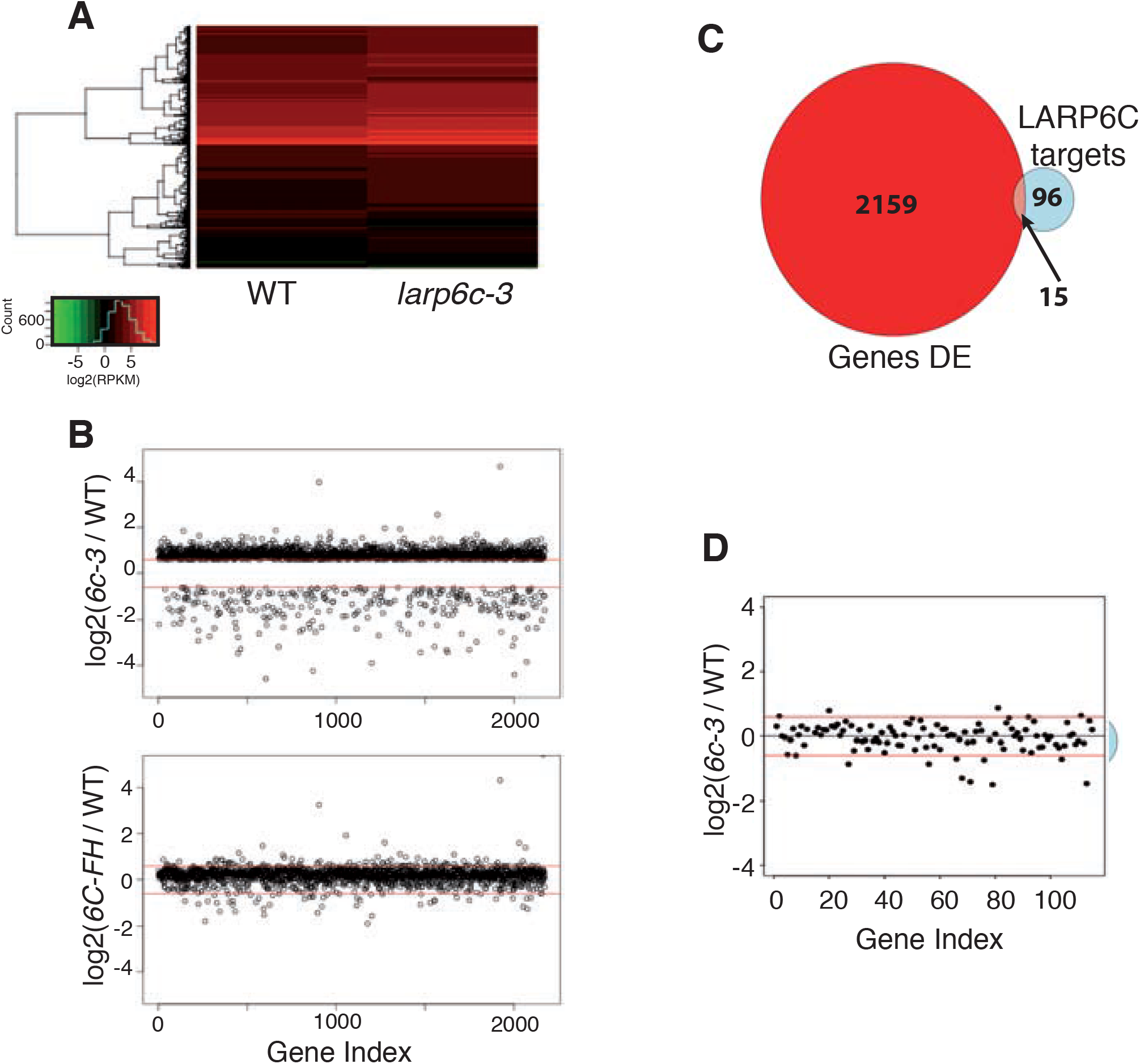
*larp6c* loss-of-function does not affect the steady-state transcripts levels of its target in dry pollen. **(A)**, Heat map representation of the log_2_(rpkm) values of genes that are differentially expressed between wild-type (WT) and *larp6c-3* plants. The heat map was built from the list of DE genes which log2(rpkm) value is between −2 and +10 (2142 genes: 98.5% of the DE genes).**(B)**, Plot representation of the log2 values of the ratios: (*larp6c-3*/wild type) (upper panel) and log2(6C-FH/wild type) (lower panel) for the 2174 genes found to be differentially expressed in *larp6c-3* mutant pollen. Red lines mark the cut off value (log2(1.5) and log2(1/1.5)). **(C)** Venn diagram representation of the number of transcripts which are DE in *larp6c-3* and/or immunoprecipitated by 6C-FH. Note that of the 115 RIP targets, 19 were not present in the transcriptomic data from the RNA-seq. **(D)** plot representation of the log2(*larp6c-3*/WT) for mRNAs identified by RIP-seq. Red lines mark the cut off value (log2(1.5) and log2(1/1.5)).

### LARP6C downregulates the translation of a reporter transcript containing a functional 5’-UTR B-box motif

To gain a mechanistic insight on the impact of LARP6C binding to its mRNA targets, we monitored the expression of a YFP reporter placed under the control of a modified 5’-UTR region belonging to a LARP6C target (At5G20410, *MGD2*) which carries a potential *bona fide* B-box and no A-box motif (Table S1). We respectively replaced the native B-box by two sequences for which we proved *in vitro* that they are strongly bound (B1-box) or not bound at all (B3-box) by LARP6C La-module (Figure 4B-E). We inserted the modified 5’-UTR sequences downstream of the CaMV35S promoter and fused to coding sequence of the YFP (*B1-YFP* and *B3-YFP*) on a plant binary vector (Figure 6A). These YFP containing plasmids were transiently expressed in *Nicotiana benthamiana* leaves individually or co-infiltrated with a plant binary vector expressing a tRFP-tagged LARP6C also under the control of the CaMV35S promoter. Three days post infiltration, leaves were collected and YFP mRNA and protein levels were assessed (Figure 6B and C). Quantitative analyses showed that when the LARP6C protein is present, the levels of the *B1-YFP* transcript are increased by two fold (Figure 6B). Conversely co-expression of LARP6C with the *B3-YFP* construct did not impact significantly the *B3-YFP* mRNA levels (Figure 6B). Remarkably, the presence of LARP6C negatively impacted the translation of the *B1-YFP* transcript significantly reducing YFP protein accumulation (Figure 6C). No altered *B3-YFP* protein levels was observed in the presence of LARP6C (Figure 6C). We also monitored the subcellular distribution of the tRFP-LARP6C fusion following agroinfiltration. Similar as observed in mature pollen grains (Figures 2, 3 and S3D) and previously in onion epidermis cells following stress exposure (Merret et al. 2013b), LARP6C formed foci in the cytoplasm of Nicotiana leaves (Figure 6D). Since in pollen grains and onion epidermis cells LARP6C foci contain the poly(A) binding protein, we reason that LARP6C likely aggregates into mRNP stress granules also in tobacco. Furthermore, the presence of LARP6C in these foci coupled with the B-Box dependent impact of LARP6C on YFP mRNA levels (Figure 6B) suggest that the higher levels observed for the B1 version results from a post-transcriptional stabilization rather than a transcriptional effect. Hence LARP6C binding downregulates its targets’ translation and protect them from decay. This mRNA protective effect of LARP6C is in contrast with our RNA-seq analyses from MPG but might be the result of distinct cellular contexts, *i*.*e* sporophytic versus gametophytic (see discussion).

**Figure 6:**
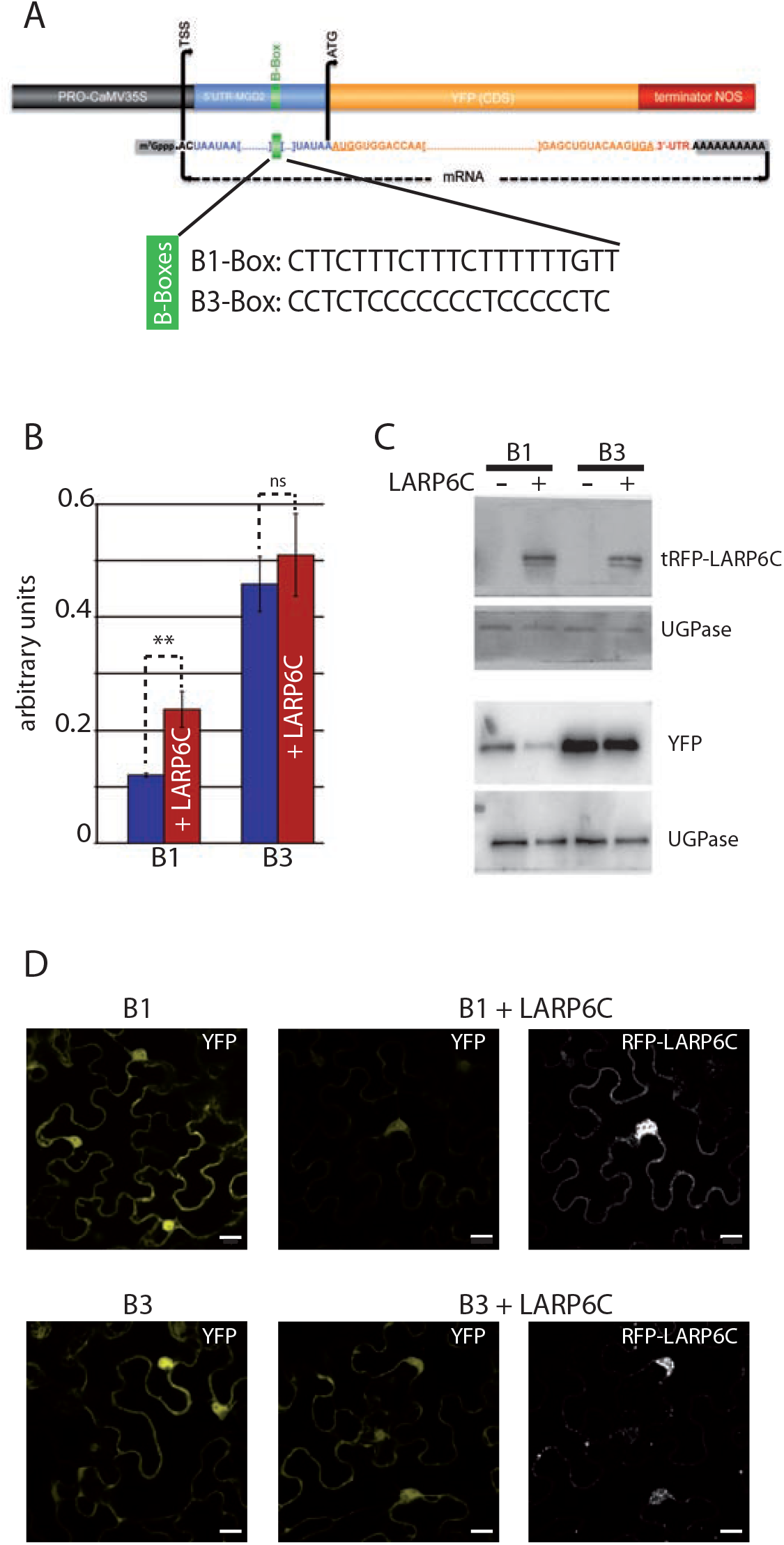
LARP6C binding at the 5’UTR reduce translation and increases mRNA levels. **(A)** Schematic representation of the YFP reporter constructs, **(B)** RT-qPCR monitoring of *YFP* mRNA levels. To normalize *YFP* mRNA levels to transformation efficiency we used the levels of the *HPTII* mRNA, encoded by the *HPTII* gene carried by the YFP binary plasmid but not the tRFP-LARP6C one. SDs were calculated from three biological replicates. p-value were obtained using a student-t test. **P < 0.005, n.s: non significant. **(C)** Western blot analyses of tRFP-LARP6C and YFP protein levels. Two western blots were prepared and respectively hybridized with anti-LARP6C or GFP antibodies. Levels of UGPase were used as loading control. Representative images of three replicates are shown. **(D)** Confocal imaging of YFP from leaves not transformed with tRFP-LARP6C (left panels) and of YFP and tRFP-LARP6C distribution (right panels). Scale bars represent 20 μM. Leaves that were observed are different from those used to prepare total RNA and protein extracts. Representative images of three biological replicates are shown.

### Partial inactivation of monogalactosyldiacyglycerol (MGDG) synthases aggravates the *larp6c* pollen tube guidance defect

Amongst LARP6C mRNA targets several, enriched in pollen, code for proteins required for lipid synthesis including MGD2, the pollen specific (our data and (Wang et al. 2008)) MGDG synthase (Botte et al., 2011). *In situ* labelling of Arabidopsis pollen tube membranes shows the presence of DGDG. And the chemical inhibition of MGD enzymatic pathway with galvestine-1 causes reduction of MGDG content and downregulates pollen tube elongation rates (Botte et al. 2011), supporting a role for MGD in male fecundation.

*MGD2*, is likely to be a direct target of LARP6C as it carries in its 5-UTR, a B-box motif with a majority of U over C residues and two stretches of three Us. We reasoned that if one of the molecular bases of *larp6c* guidance defect relates to the misregulation of MGD2 protein expression or translation, the specific inhibition of its enzymatic activity with galvestine-1 should alleviate or aggravate the *larp6c* lof phenotype. We hence monitored pollen tube elongation rate and pollen tube guidance in the presence of galvestine-1 by SIV assays (Figure 7). As previously reported (Botte et al. 2011), we observed that galvestine-1 application equally reduced pollen tube elongation of wild-type and *larp6c-3* mutant pollen (Figure 7A). However, when we monitored pollen tube guidance efficiency, wild-type pollen tubes were only slightly or not affected at all, whereas pollen tubes of *larp6c* mutants showed considerable decrease in ovule targeting in the presence of galvestine-1 (Figure 7B). In these conditions, only 30 to 60% of ovules were targeted by *larp6c*-pollen tubes whereas the mock treated samples showed 60 to 70% targeting. In some pistil explants, *larp6c* pollen tubes showed no ovule targeting at all in the presence of MGD inhibitor, a situation never observed with *larp6c-3* pollen tubes in the absence of the MGD inhibitor (Figure 1D, 7B). Collectively, these results support a synergic effect between the loss of LARP6C and the pharmacological inhibition of MGD2 synthase activity on pollen tube guidance but not on pollen tube elongation.

**Figure 7:**
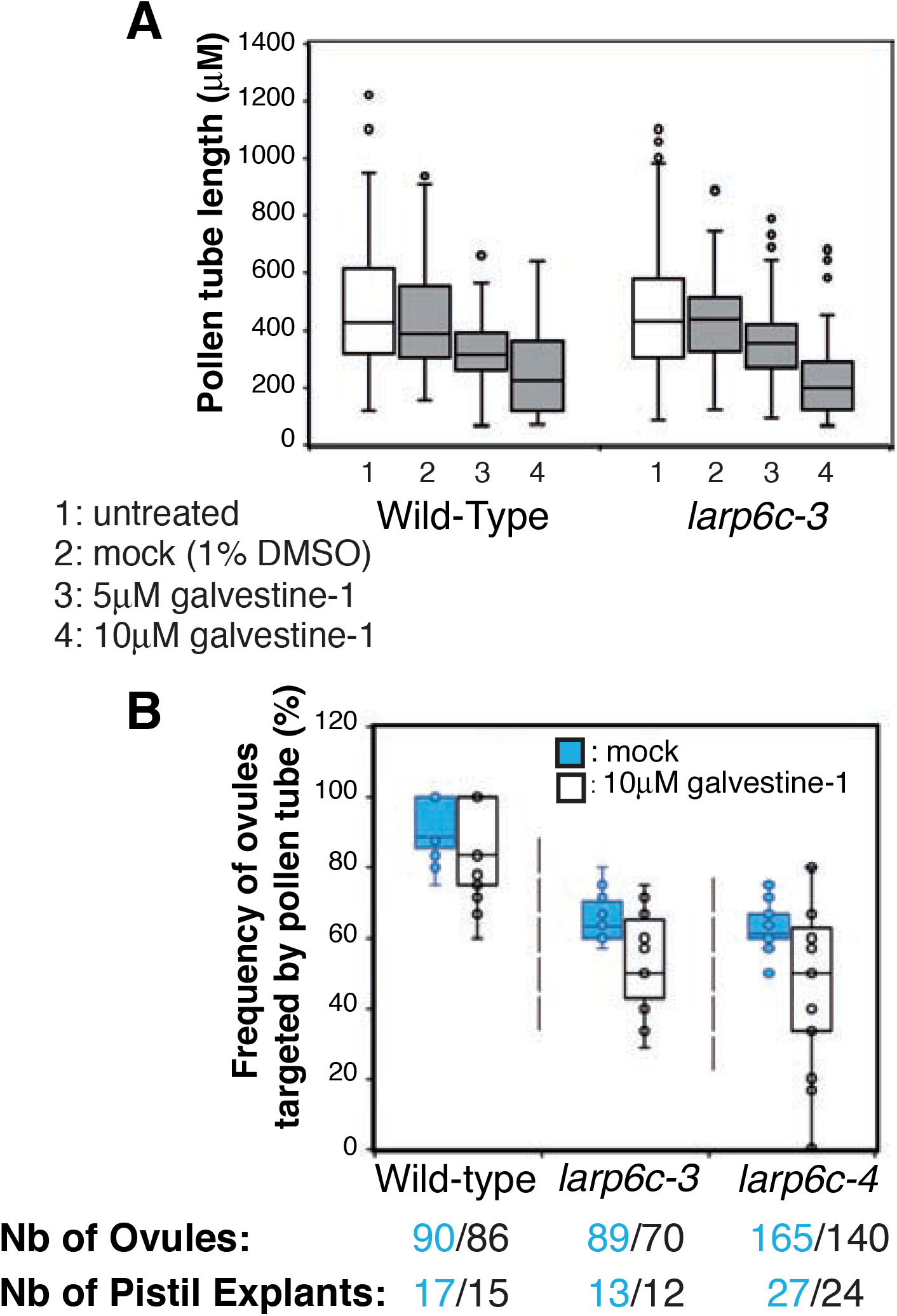
The pollen tube guidance defect of *larp6c* lof mutants is aggravated in the presence of galvestine-1. (**A**), Monitoring of pollen tube growth in the presence of galvestine-1. Whisker boxplot representation of the length of wild-type or *larp6c-3* pollen tubes grown *in vitro* in the presence or absence of galvestine-1. (**B**), Semi *in vivo* pollen tube guidance assays. Assays were conducted either in the presence of 1% DMSO (untreated) or in the presence of 10 μM of galvestine-1. Results are represented as whisker boxplots.

## DISCUSSION

We report here that the RNA binding protein LARP6C is essential for male fertility during the fertilization process at the pollen tube guidance step. Considering that SIV and *in vivo* assays showed an identical penetrance of *larp6c* pollen tube guidance defect and since the transmitting tract is absent in semi *in vivo* approaches, we conclude that LARP6C is likely to be involved in the micropylar guidance step. In other words, LARP6C is likely required for pollen tubes to sense and/or respond to ovule emitted cues.

In our study, we identified 115 mRNAs that in mature pollen, are in a complex with LARP6C. RNA-protein binding assays demonstrated that the La-module of LARP6C is able to interact with the B-type motifs that carry a majority of U over C residues and/or stretches of three consecutive Us. This is consistent with our previous observation that LARP6C shows a strong preference for oligo(U) stretches and little or no binding to any other type of homopolymers (Merret et al. 2013b). We hence hypothesize that *in vivo* LARP6C, through its La-module, interacts directly with transcripts harbouring B-boxes of the B1 or B2 type in their 5’-UTRs (*i*.*e* 52 of the 115 transcripts identified by RIP-seq (Table S1)). At this stage we cannot exclude the possibility that LARP6C also directly binds to its other RIP targets through A-boxes, perhaps using yet unidentified RNA binding regions outside the La-module. Or that it is in a complex with these other RIP-identified mRNAs but not through direct binding. We propose that the post-transcriptional regulations mediated by LARP6C on the RIP-identified mRNAs in mature pollen are necessary for ovular guidance of the pollen tube. Indeed a gene to gene analysis of the list of LARP6C targets allowed us to recognize genes which functions were directly found to be necessary for pollen tube guidance, polarized tip growth, signalling or membrane synthesis. Considering the role of LARP6C in male fecundation, we anticipate that amongst its target genes several are yet unrecognized actors of the guided growth of the pollen tube. As an example, our work provides evidence that MGD2 is a novel actor of this cellular process. But then, what is the mechanistic impact of LARP6C binding to the 5’-UTR of its target transcripts? In eukaryotic cells, mRNAs bound by various proteins tend to aggregate in the cytoplasm into self-assembling membraneless structures to form so called mRNP granules. Several types of mRNP granules, defined upon their cellular context, presumed functions and the presence of particular protein markers have been identified and classified (Buchan 2014). But one of their common features is that they all contain repressed mRNAs that are capable of (re)-entering translation following appropriate signals. The use of a transient reporter system in tobacco leaf cells allowed us to observe that the presence of LARP6C upregulates the mRNA levels, likely through increase in mRNA stability, of its targets while antagonizing their translation. In these transient assays LARP6C systematically displayed a punctuate distribution in the cytoplasm that we deem likely to be stress granules. In somatic cells, stress granules are known to protect translationally silent mRNPs that need to be swifty re-introduced in translation after the stress ends (Merret et al. 2017). We propose that the binding of LARP6C at the 5’-UTR acts to repress translation of its mRNA targets and promote their accumulation into aggregates as storage granules and protect them until their translation is reactivated (see model in Figure 8).

**Figure 8:**
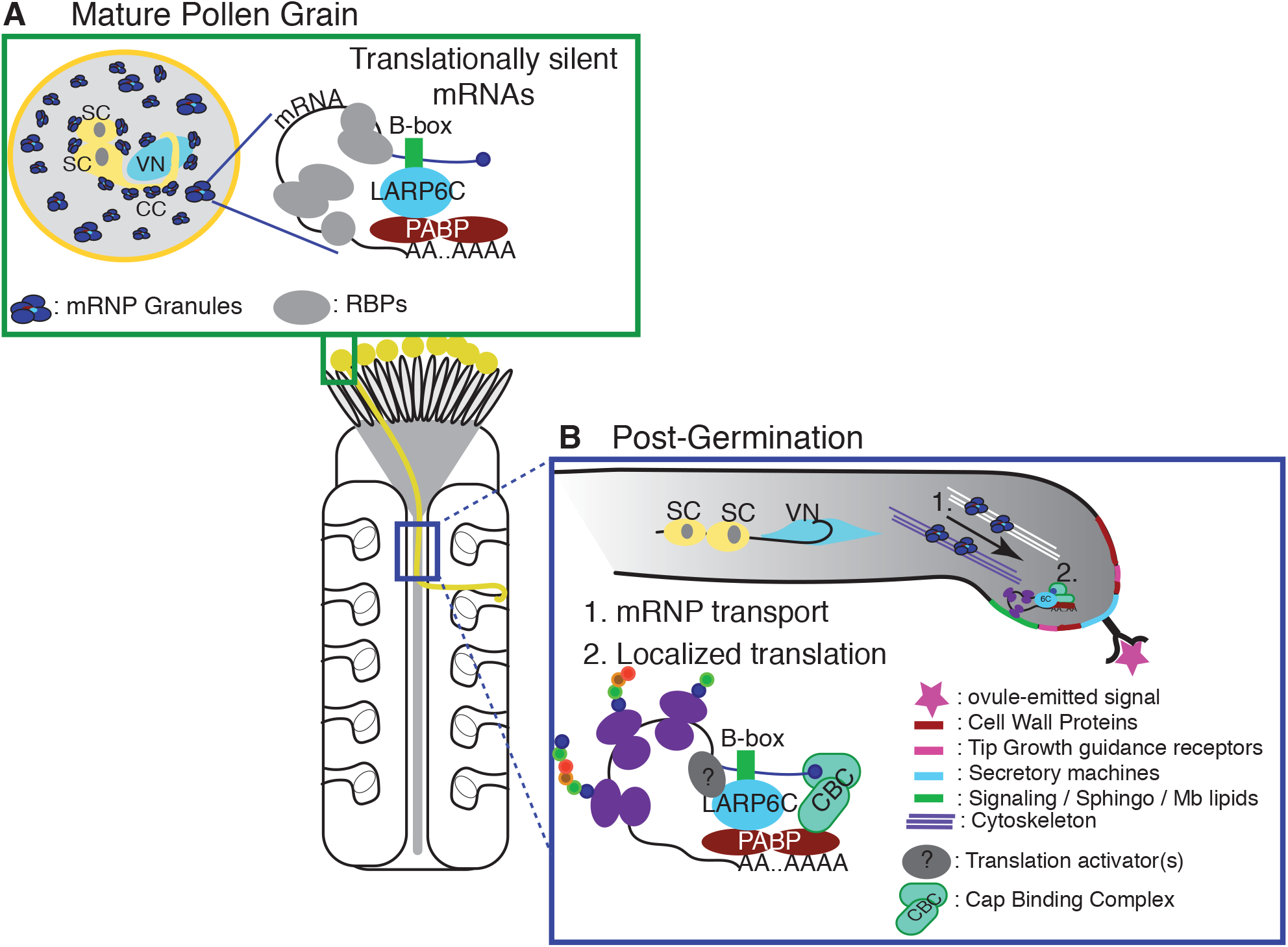
A model of LARP6C-B-box-PABP mediated translational silencing in the pollen tube guidance process. **(A)** In dry pollen grain, LARP6C binds to B-box 5’UTR motif of its mRNA targets and together with the poly(A) Binding Protein (PABP) localises to cytoplasmic aggregates to maintain its targets mRNA in translationally silent state. **(B)** During pollen tube tip growth, LARP6C could accompany its bound mRNA targets (as shown in human LARP6) and keep them translationally repressed until an external signal (likely ovule-emitted cue) is perceived at the pollen tube tip. Perceived paracrine signal would then relieve LARP6C-mediated repression through a yet unknown process to permit localized translation of factors involved in guided pollen tube growth including localized secretory machinery, receptors and membrane lipids. This is consistent with the observation that the chemical inhibition of MGD2 enzymatic activity aggravates guidance defect but not pollen tube growth.

We next asked whether LARP6C retains such molecular functions in the non somatic cells *i*.*e* MPG and pollen tube. Interestingly, LARP6C forms punctuate foci throughout the vegetative cell cytoplasm and appears to accumulate, also with an aggregate pattern, around the sperm cells and vegetative nucleus. These LARP6C foci fully co-localized with the poly(A) binding protein PAB5 supporting the idea that these aggregates are likely mRNP granules. Scarpin et al. (Scarpin et al. 2017) reported that the *SK14* mRNA which was found to be present but translationally silent in Arabidopsis and Tobacco MPGs and reactivated at germination (Honys et al. 2000; Scarpin et al. 2017), forms punctuate foci that also contain DCP1 and VARICOSE that are components of p-bodies in somatic cells (Weber et al. 2008). Interestingly, at germination the production of the SK14 protein coincides with the disappearance of *SK14* foci, suggesting that mRNAs could be stored until their protein product is required at or along the progamic phase (Scarpin et al. 2017). We hence propose that LARP6C in dry pollen, could help maintaining its targets in aggregates in a translationally silent form until their translation product is required for the pollen tube function.

The stabilization effect of LARP6C binding observed in transient tobacco assays is in contrast with the observation that in pollen grains LARP6C does not seem to control its targets’ steady-state levels. In pollen, as in animal gametes, mRNAs ((including LARP6C target transcripts in *larp6c* lof pollen) were found to be highly stable suggesting that in the male gametophyte the mRNA decay machineries, as translation, are poorly active (Ylstra and McCormick 1999). This assumption is consistent with our observation that DCP1 which is a core constituent of the decapping mRNA turnover complex, localises into foci in mature pollen grains representing quenched DCP1 activities. Indeed, work conducted in animal cells proposes that when components of the mRNA decay machineries are present in p-bodies they are catalytically inactive (Hubstenberger et al. 2017). Therefore, the discrepancies between results from transient assays in tobacco leaf cells and RNA-seq from pollen grains could hence be the consequence of differences in how mRNA decay complex operates between sporophytic versus reproductive cells as exemplified in other cellular machineries such as Transamidase GPI8 GPI-anchoring complex (Liu et al. 2016).

Using a specific chemical inhibitor of MGD2 synthase activity we observed an aggravation of *larp6c* pollen guidance. This observation suggests that loosing LARP6C-mediated post-transcriptional regulations and downregulating MGDG/DGDG production add up to dampen further pollen tube competence. Hence this suggests that, in pollen tube, LARP6C might stabilize *MGD2* mRNAs and/or promote their translation rather than inhibit protein production. But to be consistent with the observation that it is the guidance defect of *larp6c* pollen tubes that is aggravated and not their growth rate in the presence of galvestine-1, we propose that LARP6C might shift to a translation stimulatory role only when it reaches together with its bound mRNA, the plasma membrane at the site of ovular cue perception to permit the synthesis of MGD2 exactly where its synthase activity is needed (see Figure 8). This working model, is consistent with the known role of human LARP6 that shifts from a repressor to an activator role in the coordinated translation of its target mRNAs, according to its associated proteins and subcellular localization ((Zhang and Stefanovic 2016)).

In mammals, the nervous system axons and dendrites need to rapidly extend unidirectionally into polarized cells guided by chemical cues. In this process, reminiscent of the guided growth of the pollen tube, neuronal mRNP transport granules play a crucial role. They act to transport translationally repressed mRNPs to the precise site of their translational activation and protein product function at the tip of an axon (Loya et al. 2010). Our work posits the exciting notion that there could exist a similar system in the process of plant fecundation.

## METHODS

### Plant material and growth conditions

*Arabidopsis thaliana* ecotype Columbia-0 (Col-0) was used as wild-type reference. The *larp6c-3* (SAIL_268E02) (McElver et al. 2001) and *larp6c-4* (WiscDsLox293-296invG4) (Woody et al. 2007) lines were respectively ordered from the NASC and ABRC stock centres. Genotyping primers are reported in Table S4. Stable transgenic lines were obtained using the *Agrobacterium tumefasciens* based floral dip technique (Clough and Bent 1998). For *in vitro* culture, surface sterilized seeds were sown on synthetic Murashige & Skoog (MS) medium at 2.20 g/L (1/2 MS) with 0.8% agar and stored for 48 h at 4°C in the dark before being grown under continuous light (50-60 □E.m^-2^.s^-1^) at 20°C. Soil cultures were performed in growth chambers with 16 h light (100□□E.m^-2^.s^-1^) / 8 h night at 20°C with approximately 75% humidity. See supplemental experimental procedures for growth conditions used to collect mature pollen for RNA-seq and RIP-seq experiments.

### Cloning and plasmids

All LARP6C containing fusions (YFP, tagRFP and FLAGHA) were expressed under the upstream genomic sequences (spanning region −1181 to −1 from ATG) of the *LARP6C* gene (AT3G19090 locus). To obtain the LARP6C-tRFP and LARP6C-FLAGHA fusions, we PCR amplified from Col-0 total genomic DNA, the *LARP6C* genomic region starting from nucleotide −1181 from the ATG to the last nucleotide before the stop codon, using primers 543 and 544 (Table S4). The PCR product was inserted at sites *KpnI* and *NheI* either upstream to the tagRFP or FLAG-HA tag into a pCAMBIA1300-based plant binary vector expressing the *HPTII* (hygromycin) resistance gene, giving rise to vectors pEB16 and pEB13. To obtain the YFP-LARP6C expressing binary vector, the genomic region spanning nucleotides −1181 to −1 from the ATG of the *LARP6C* gene was PCR amplified with primers 974 and 975 from vector pEB13 and cloned at sites *SacI* and *KpnI* upstream to a gateway entry cassette into a pPZP221 (Hajdukiewicz et al. 1994) based plant binary vector, carrying the *AAC1* (gentamycin) resistance gene, giving rise to vector p788. The *LARP6C* coding genomic region was PCR amplified from total genomic DNA with primers 976-977 and cloned at sites *Spe*I-*EcoR*V downstream to the YFP tag located between the AttL1 and L2 gateway donor sites, giving rise to plasmid p793. The plant binary vector, p795, was obtained through a gateway cloning reaction between p788 and 793. The *PAB5* genomic sequence (expanding from −1966 from the ATG to the last nucleotide before the stop codon) was PCR amplified from total genomic DNA with primers eb3 & eb4 and cloned using the restriction endonucleases *Xba*I and *BamH*I into vector CTL579 (pCAMBIA-1300) upstream to the GFP tag, giving rise to plasmid pEB3. The binary vector pGEX2-GEX2GFP was obtained from (Mori et al. 2014). Binary plasmids for transient assays were prepared as follow. DNA fragments encompassing a basal CaMV35S promoter without enhancer sequences (Topfer et al. 1987), modified *MGD2* 5’-UTRs and the YFP CDS were ordered from GeneCust (http://www.genecust.com/fr) and subcloned into a gateway donor vector. DNA casettes were then introduced into a plant binary vector containing the *HPTII* (hygromycin) resistance gene (vector CTL575, a derivative from pCAMBIA-1300). The *LARP6C* coding sequence was fused downstream to the tagRFP fluorescent reporter and placed under the control of a strong CaMV35S promoter by gateway recombinaison with the plant binary entry vector p-SITE-6C1 (Martin et al. 2009).

### Pollen phenotyping

Pollen transmission tests were conducted through hand pollination of wild-type pistils with pollen from mutant and complemented homozygous plants. The number of offsprings carrying the mutation was scored either by PCR-based genotyping or by selection of antibiotic/herbicide-resistant seedlings. The number of mutant seedlings was expressed as a percentage over the total number of seedlings and transmission efficiency of the mutant gametophyte calculated as TE= ((No of mutant / No of WT) x 100). Experiments were conducted in triplicates. Pollen tube guidance efficiency was scored either through semi *in vivo* (Palanivelu et al., 2006) or *in vivo* assays by observation of pistils hand pollinated with the mutant pollen of interest and stained with aniline blue (Mori et al., 2006). Emasculated pistils were from *ms1* (male sterility 1) heterozygous plants. This garantees that pistils were not contaminated with wild-type pollen as the *ms1-1* mutant is male sterile (Van der Veen and Wirtz 1968; Ito et al. 2007). Monitoring of pollen tube guidance in the presence of galvestine-1 was conducted through semi *in vivo* assays but on plates containing 1% DMSO for mock treatment or 5 or 10 μM of galvestine-1 with a final concentration of 1% DMSO.

### Confocal microscopy

Subcellular localization experiments were conducted with an LSM700 confocal microscope (Zeiss) using emission/excitation wavelengths: 405 nM / 420-480 nM for DAPI, 555 nM / 600-700 nM for tRFP and 488 nM / 490-555 nM for YFP and GFP. Mature pollen grains were collected through dipping of open flowers into a DAPI solution (1X PBS pH 7, 0.5% triton-X100, 1 □g/mL 4’,6-diamino-2-phenylindole) drop deposited on a glass slide. To collect immature pollen grains from microspore, bi- and tricellular stages, anthers were collected from unopened buds, placed in a DAPI solution drop on a glass slide and gently crushed to free pollen grains from pollen sacs. Pollen tubes germinated and grown as described in the supplemental experimental procedures were directly observed from the germination slide.

### RNA related techniques

Total RNA extractions were performed according to (Logemann et al. 1987). To purify total RNA from mature pollen, 0.5 to 3 mg of dry pollen were grinded with a Silamat S6 (Ivolcar Vivadent, Schaan, Liechtenstein) in 500 □L of GuHCl buffer (8 M GuHCl, 20 mM MES, 20 mM EDTA, at pH 7, 50 mM β-mercaptoethanol) and with 5 glass beads. Following 1 min vortexing at full speed and 10 min incubation on ice, RNAs were separated from proteins with two consecutive phenol-chloroform-IsoAmylAlcool (IAA) (25:24:1) extractions, followed by one extraction with chloroform-IAA (24:1), consisting in 1 min vortexing at full speed and 10 min centrifugation at 16°C, 14.000 g. RNAs are then precipitated in the presence of 0.7 vol isopropanol, 1/20^th^ CH_3_COOH at 1 M for 20 min at −20°C. Following 20 min centrifugation at 4°C, 14.000 g the RNA pellet is washed in 90% EtOH and resuspended in RNAse free water. The average efficiency of pollen RNA extraction was at 5-6 μg of RNA per mg of dry pollen.

RNAs were treated with DNAse using the Turbo^™^ DNAse kit from Ambion (Thermofisher), following the supplier instructions for the “Rigourous DNAse treatment” procedure. For Reverse transcription, 500 ng of DNAse treated RNAs were reverse transcribed with the SuperScript III reverse transcriptase (Life Technologies, Thermo Scientific, USA) and 500 ng of oligodT_18_. PCR reactions were conducted with 1.5 μL (1/20^th^ of the reverse transcription solution) of cDNA for LARP6C and 2 μL for ACTIN8 amplication, using the GoTaq G2 Flexi DNA polymerase from Promega and 30 PCR cycles. For qPCR analyses, total RNA was isolated using the Monarch Total RNA Miniprep Kit (New England Biolabs), and genomic DNA were eliminated by gDNA removal columns and on-column DNaseI-treated (New England Biolabs). Following, 5 μg of total RNA were reverse transcribed with the SuperScript IV kit (Life Technology) using an oligodT_18_ primer. qPCR reactions were conducted on the LightCycler 480 Multiwell Plates 384-well themocycler (Roche Applied Sciences), with the following amplification program: 5 min 95°C, 40 cycles of 15 s at 95°C, and 1 min at 60°C. After the amplification, a melting curve analysis with a temperature gradient of 0.1°C/s from 60°C to 95°C was performed. The PCR mix contained Takyon qPCR master mix (Eurogentec), 500 nM gene-specific primers, and 1 μL cDNA in a total reaction volume of 10 μL. Primer efficiencies were determined on standard curves, with 10-fold serial dilution series, ranging from 1 × 10^1^ to 1 × 10^6^ of the cDNA samples. Relative quantification was performed by the ΔΔC_T_ method (Livak and Schmittgen 2001), with *YFP* and *HPTII* genes used as the target and reference genes, respectively. The sequences of the primers used for RT- and qPCRs are reported in Table S4.

### RNA-seq, RIP-seq, bioinformatics

Techniques related to large scale analyses (RNA-seq and RIP-seq), as well as the bioinformatic analyses are detailed in the supplemental experimental procedure section.

### Protein related techniques

Total protein extracts were separated by SDS PAGE on an acrylamide gel before being electrotransferred on a PVDF membrane (0.45 μM, Merck millipore). After saturation in a (1X TBS −0.1% tween - 5% fat-free dry milk) solution, membranes were incubated with the appropriate primary then secondary antibodies, washed three times with a 1X TBS −0.1% tween solution, and chemiluminescence revealed with the EMD Millipore immobilon western chemiluminescent HRP substrate kit. Antibodies against the LARP6C protein were custom made (at Agro-Bio, La Ferté St Aubin, France) through immunization of rabbits with peptide “KRTSQFTDRDREELGQ” (amino acids 220-235). Anti-LARP6C antibodies are utilized at 1/1000^th^ dilution in a (1X TBS −0.1% Tween - 5% fat-free dry milk) solution and incubated overnight at 4°C with gentle shaking. Secondary antibodies were anti-rabbit HRP (Bio-Rad) and used at a 1/5,000^th^ dilution. Anti-ACTIN antibodies were from Affinity Bio Reagents (Golden, CO, USA) and used at 1/15,000^th^ dilution. Secondary antibodies were anti-mouse HRP (Bio-Rad) used at a 1/10,000^th^ dilution. For transient assays, the YFP protein was detected using the GFP antibody (Clontech) at 1/5,000^th^ and the UGPase antibody (Agrisera) at 1/7,500^th^ dilution, both incubated overnight at 4°C. Secondary antibodies anti-mouse HRP (Bio-Rad) used at a 1/2,000^th^ dilution, and anti-rabbit HRP (Bio-Rad) used at a 1/5,000^th^ dilution, were incubated, respectively for YFP and UGPase, for 1h at room temperature.

The LARP6C La-module (amino acids 137-223) was recombinantly expressed as in (Merret et al. 2013b). The hexahistidine tag (His-tag) was removed by proteolysis through incubation at 4°C overnight with HRV 3C Protease in 50 mM Tris-HCl pH 7.25, 100 mM KCl, 0.2 mM EDTA, 1 mM DTT, 10% gycerol buffer. The cleaved His-Tag and the protease were separated from the La-module through retention onto a Ni-NTA column. The protein was then subjected to heparin column purification and dialysis in a final buffer as in (Merret et al. 2013b).

### ITC and EMSA

RNA oligos were ordered as custom made from Iba-Lifescience GmbH (Göttingen, Germany) at a 1 μMolar scale and HPLC purified. The lyophilized RNAs were resuspended in RNAse free water (DEPC treated) at concentrations ranging from 2.5 to 3 mM. The ITC experiments were performed as in (Merret et al. 2013b) on a iTC200 microcalorimenter (Malvern). Here, 20 injections of 2 μL RNA solutions at 200-320 μM concentration were added to protein solutions (6C-La-module) at 20-30 μM with a computer-controlled 40 μL syringe. Control titrations of RNA into buffer alone and into protein solution without the His-tag were performed (not shown). The heat per injection normalized per mole of injectant versus molecular ratio, was analyzed with the MicroCal-Origin 7.0 software package and fitted using a nonlinear least-square minimization algorithm using a theoretical single-site binding model. ΔH (reaction enthalpy change in kcal/mol), K_b_ (binding constant equal to 1/Kd), and n (molar ratio between the two proteins in the complex) were the fitting parameters. The reaction entropy was calculated using the relationships ΔG =-RTln K_b_ (R = 1.986 cal/mol K; T = 298 K) and ΔG = ΔH -TΔS. All ITC experiments were repeated at least three times, and a detailed analysis of the thermodynamic parameters and errors calculated as the standard deviation from the mean value are reported in Table S2. For EMSA (Electrophoretic Mobility Shift Assay), RNAs were 5’-labelled with γ-^32^P-ATP using the T4 polynucleotide kinase. Recombinant protein and RNA were mixed and incubated for 15 minutes at room temperature in (20 mM Tris pH 7.25, 200 mM KCl, 5% of glycerol, 1 mM DTT, 0.1 mg/mL of BSA and 0.7 units of RNAse inhibitors (RNAseOUT, Invitrogen)). Each reaction mixture (22 μL) contained 3 nM of labelled RNA oligo and varying amounts of LARP6C La-module (3-fold serial dilutions from a 88 μM sample). The experiments were performed in the absence or in the presence of unlabelled tRNAmix of *E. coli* MRE 600 (0.01 mg/mL). After the addition of 2 μL of 30% Ficoll, the samples were loaded on a 9% native polyacrylamide gel prerun at 100 mV for 1 hour at 4°C in 0.5x TBE (Tris-borate-EDTA buffer). A typical EMSA experiment was run at 4°C for 1 hour at 125 mV. Gels were dried onto 3MM chromatography paper and then exposed to a phosphoimaging plate overnight. These were then analysed on a phosphoimager Typhoon Trio and quantified with Image Quant TL software. The fraction of bound RNA was plotted versus the protein concentrations. To determine the dissociation constants the data were fitted with Origin 8.0 to a modified Hill equation (Ryder et al. 2008) using non-linear least squares methods and assuming a 1:1 stoichiometry.

### *Nicotiana benthamiana* transient assays

Plasmids B1-YFP, B3-YFP and RFP-LARP6C were respectively transformed into *Agrobacterium tumefasciens* strain LB4404. Transformed bacteria were cultivated until they reached OD 0.8 and used to infiltrate fully expanded leaves from eight-week-old *N*.*benthamina* plants, using needless syringe, as described in (Ruiz et al. 1998). When double transformations were performed (B1-YFP and RFP-LARP6C or B3-YFP and RFP-LARP6C) an equimolar mixture of Agrobacterium cultures (OD 0.4) was used for transformation. For each transformation we also used an Agrobacterium culture carrying a plasmid (OD 0.2) allowing the expression of the P19 suppressor of RNA silencing (Voinnet et al. 2003). Three days post-infiltration (3 dpi), transformed leaf segments were collected, flash frozen and total RNA or protein extracted. Alternatively, 3 dpi, transformed leaves were observed by confocal microscopy to monitor YFP and RFP signals.

### Accession Numbers

Sequence data in this study can be found under the following accession numbers : Bioproject PRJNA557669. RNA-Seq-Col0-Rep 1: SRR9887494; RNA-Seq-Col0-Rep 1: SRR9887493; RNA-Seq-*larp6c-3*-Rep 1; SRR9887496; RNA-Seq-*larp6c-3*-Rep 2 SRR9887495; RIP-Col0-Input-Rep 1: SRR9887498; RIP-Col0-Eluate-Rep 1: SRR9887500; RIP-Col0-Input-Rep 2: SRR9887497; RIP-Col0-Eluate-Rep 2: SRR9887499; RIP-LARP6C-Input-Rep 1: SRR9887490; RIP-LARP6C-Eluate-Rep 1: SRR9887492; RIP-LARP6C-Input-Rep 2: SRR9887489; RIP-LARP6C-Eluate-Rep 2 : SRR9887491.

## Supplemental data

**Table S1:** RIP-seq experiment: rpkm values for genes detected in the input and eluate fractions of wild type and 6C-FH pollen grain. List of putative LARP6C mRNA targets with a description of the two 5’-UTR enriched motifs.

**Table S2:** Thermodynamic parameters to the calorimetric analyses shown in Figure 4B, 4D and S6A.

**Table S3:** RNA-Seq experiment: rpkm values for genes detected in wild type and *larp6c-3* mature pollen grain. List of genes differentially expressed between *larp6c-3* and wild type.

**Table S4:** Primers used for genotyping, cloning and RT-PCR. Sequence, efficiency of qPCR primers, amplicon sizes and Tms.

**Supplemental Figure 1:** Expression profiles of *LARP6A, 6B* and *6C* mRNAs across development. Supports Figure 1A.

**Supplemental Figure 2:** Pollen maturation, germination and pollen tube growth of *larp6c* lof mutants. Supports Figure 1D-E.

**Supplemental Figure 3:** Schematic representation of *LARP6C* transgenes and western blotting analyses of their accumulation. Subcellular distribution of LARP6C tagged with tagRFP at its C-terminus in bicellular, tricellular and mature pollen grain. Supports Figures 1D-E, 2 and 3.

**Supplemental Figure 4:** Pollen attraction competence of *larp6c-*3 and *6c4* ovules. Supports Figure 1C-E.

**Supplemental Figure 5**: Reproducibility of the RIP-seq data, RIP-seq filtering workflow, representation of the position of A and/or B boxes on the 5’-UTRs of LARP6C targets. Supports Figure 4A-B.

**Supplemental Figure 6:** ITC and EMSA assessment of LARP6C La-module binding to A type oligos. Supports Figure 4C-F.

**Supplemental Figure 7:** Reproducibility of the RNA-seq data. Supports Figure 5.

## Supplemental text

Supplemental figure legends and supplemental methods.

## Acknowledgements

This work was supported by the CNRS, the University of Perpignan (UPVD), the Institut Universitaire de France (IUF) and the Bio-Environnement platform through utilization of the confocal microscope. This study is set within the framework of the “Laboratoires d’Excellence (LABEX)” TULIP (ANR-10-LABX-41). EB was the recipient of a PhD grant from the UPVD, Doctoral School ED305. CGL jr. is the recipient of a short-term contract supported by the ANR Heat-EpiRNA (ANR-17-CE20-007-01). SH, KK and DH are supported by GACR grant numbers 17-23203S and 18-02448S. ICG was supported by a Royal Society Newton International fellowship (ref. NF140482). IGC and MRC thank the Centre for Biomolecular Spectroscopy at King’s College London funded by a capital award from the Wellcome Trust. We would like to thank E. Maréchal (LPCV, Grenoble, France) for kindly providing us galvestine-1 and R. Merret (LGDP, Perpignan, France) fo critical reading of the manuscript.

