## Supplemental Data for "LARP6C regulates selective mRNA translation to promote pollen tube guidance in *Arabidopsis thaliana*"

**SUPPLEMENTAL DATA METHODS**

**Growth conditions and pollen collection technique for RNA-Seq and RIP-Seq assays**

RNA-seq and RIP-seq mature pollen material were obtained from plants sown, grown and harvested under the exact same conditions. Wild type (Col0), *larp6c-3* and (*larp6c-3*; PRO$^{6C}$-LARP6CFLAGHA) (abbreviated to 6C-FH) were sown on 1/2 MS - 0.8% agar in duplicate and cultivated for 15 days *in vitro* at 20°C, with 24 h light (50-60 $mE^{-1}m^{-2}$-$s^{-1}$). Seedlings were then transferred to *Seinernema feltiae* (at 5000 / soil Liter) treated soil and cultivated in a greenhouse with approximately 16 h day at 22°C night and day (greenhouse conditions, july 2013 Gif/Yvette France) for 5 weeks. Wild type Col0, *larp6c-3* and 6C-FH (transformant number 267 (see Figure S3C for a western blot analysis of LARP6C-FLAGHA protein accumulation)) plants were grown together in the same compartment of the greenhouse in large plates each containing 96 plants. Two sets of (Col0, *larp6c-3*, 6C-FH) plants were grown to produce two replicates. For each replicate 15 Col0, 3 *larp6c-3* and 15 6C-FH plates were prepared. The plates of each replicate were respectively installed on two distinct growth shelves inside a same compartment and each genotype interspersed inside each replicate. To avoid cooling air efflux and temperature gradient effects and obtain an homogeneous growth between genotypes inside each replicate, the plates were rotated along their respective shelf every 2 weeks.

Mature pollen grains were collected using a homemade pollen collector as described in (Johnson-Brousseau and McCormick 2004). Briefly, mature pollen is collected by aspiration with a vacuum cleaner equipped with three successive filters of 140, 60 and 10 μm diameter. Pollen grains are scrapped from the 10 μm filter into an Eppendorf tube, weighted, flash frozen in liquid nitrogen and stored at -80°C.

**RNA sequencing**

Total RNAs were extracted from 5 mg of wild-type and *larp6c-3* mature pollen grains as described in the methods section of the main text. RNA quality was verified by gel electrophoresis and staining with Gel-red (Biotium). DNase treatment was conducted as for RT-PCR assays, reaction stopped and DNase extracted with phenol/chloroform/IAA before RNA is precipitated and resuspended in RNase-free water. DNase treated RNA qualities were controlled with a Bio-Analyser. Libraries preparation (with a True seq Stranded mRNA kit from Illumina) and sequencing were subcontracted to Fasteris (Switzerland). Single-end (101bp read length) sequencing was performed on HiSeq 2000 with a depth of 31 to 60 M reads. Experiments were conducted in duplicate for each genotype.

**RNA Immunoprecipitation and sequencing (RIP-Seq)**

After pollen collection, crude extracts (Inputs) were prepared from identical amounts of Col0 and 6C-FH expressing pollen. Twenty-five mg of mature pollen were resuspended in 250 volumes of extraction buffer (50 mM Tris HCl pH 7.4, 10% glycerol, 100 mM NaCl, 5 mM $MgCl_2$, 80 units / ml RNAsin® Plus RNAse Inhibitor (Promega), 1X Mg132, 1% protease inhibitor cocktail for plant extract (Sigma)) and ground using a Silamat S6 before extracts are cleared by centrifugation (4°C, 14.000 g for 10 min). 700 and 40 μL of Input fraction were saved to purify total RNA (Input RNA) and total protein (Input Protein) to respectively conduct RNA-sequencing and western blotting. For each sample (Col0-Replicate 1, Col0-Replicate 2, 6C-FH-Replicate 1 and 6C-FH-Replicate 2), the immunoprecipitation step was conducted with 5 mL of crude extract. For each sample 250 μL of Anti-Flag M2 (Sigma) magnetic beads were thoroughly washed with 10 volumes of extraction buffer and evenly divided between the 5 tubes. After removing the supernatant, 1 mL of crude extract (approximately 300 μg of total protein) were added to each tube and rotated for 1 h at 4°C, on a wheel at 10 rpms. The unbound fractions were collected, pooled and 700 and 60 μL stored for RNA and protein analyses. Beads were washed once with 5 vol of washing buffer (50 mM Tris HCl pH 7.4, 10 % Glycérol, 150 mM NaCl, 3 mM $MgCl_2$, 80 units / ml RNAsin® Plus RNAse Inhibitor (Promega), 1X Mg132, 1% protease inhibitor cocktail for plant extract (Sigma), 0.1% Triton™ (Sigma)) before being transferred to a new tube, being pooled and washed twice more in 2 vol of washing buffer. The elution step was conducted as follow: 6% of the beads (15 μL) were resuspended in 1X Laemli buffer, heated for 10 min at 100°C and the supernatant used to run a western blot analysis to monitor the efficiency of immunoprecipitation of the 6C-FH protein. The remaining 94% of the beads were resuspended into 400 μL of guanidium hydrochloride buffer, incubated for 10 min at RT, the supernatant collected and separated from proteins with two phenol/chloroform/IAA extractions before the RNA is precipitated in the presence of

80 μg of glycogen and resuspended into 10 μL of RNase-free water. RNAs in the eluate fractions were quantified with a nanodrop and their quality varified with a Bio-Analyzer system. The library preparation and sequencing (Input RNA and Eluate RNA) were subcontracted to Fasteris (Switzerland). The libraries constructed with the input RNAs were prepared from poly(A) purified RNAs using the True seq Stranded mRNA kit from Illumina and the libraries prepared from the eluate fractions were constructed from unpurified RNAs. Sequencings were performed as single-end 101 bp and with a depth of 42 to 64 M reads for the inputs and 25 to 40 M reads for the eluates.

**Bioinformatics**

For each RNA-seq and RIP-seq sample and replicate, read quality was firstly verified with the FastQCsoftware (http://www.bioinformatics.babraham.ac.uk/projects.fastqc) and clusters with a Qscore below 30 filtered out. 95-96% of the RNA-seq clusters and 92-94% of the RIP-seq clusters had a Qscore of at least 30 (please see Tables S1 and S3 for the sequencing depth and quality scores of each sample). In a second step non nuclear genomic sequences and ribosomal RNA sequences were filtered out against the TAIR10 database using Bowtie2 (Langmead and Salzberg 2012) and the remaining clusters aligned against TAIR10 (TAIR10_genes_transposons.gtf) with TopHat2 (Kim et al. 2013). After filtering, more than 99% of the clusters were retained for the RNA-seq samples and inputs of the RIP-seq. For the eluates, 5% of the Col0 clusters and 15% of the 6C-FH clusters were retained, because of the high contamination with ribosomal RNAs. Normalized mean values were attributed to each gene, and differentially expressed genes identified with the Cufflinks, Cuffmerge and Cuffdiff suite (Trapnell et al. 2010). For the RNA-seq and RIP-seq assays, genes that did not display at least 1 rpkm (read per kilobase per million mapped reads) value in one of the conditions, were filtered out. To search for a motif shared by putative targets of LARP6C we firstly looked at the Araport database (https://www.araport.org/) for their 5' and 3'-UTR sequences and retrieved sequences for 112 out of 115 putative targets. We then ran MEME searches (at the MEME suite portal (http://meme-suite.org/)) using, as control sequences, a set of 112 randomly chosen 5'- and 3'-UTR sequences from genes expressed in pollen according to our transcriptomic analysis.

**Pollen phenotyping : pollen maturation, pollen tube germination and growth**

To monitor pollen maturation defects, mature pollen were collected from wild-type and *larp6c* homozygous plants, deposited on a DAPI solution (1X PBS pH7, 0.5% Triton, 1□g/mL 4',6-diamidino-2-phenylindole (DAPI)) and the number of mutant pollen with undivided generative cells were determined using confocal microscopy. Pollen tube germination and growth defects were monitored *in vitro* or through semi *in vivo* assays. To grow pollen tubes *in vitro*, we used a modified version of the protocol from (Boavida and McCormick 2007). The pollen tube growth medium (0.01% boric acid, 5 mM $CaCl_2$, 5 mM KCl, 1 mM $MgSO_4$, 10% sucrose, pH 7.5 (KOH), 1.5% phytagel) is prepared extemporaneously and directly poured onto a microscope slide. Mature pollen is deposited on the solid medium and slides incubated for 8 h in a humid chamber at 20-22°C in the dark. Pollen tubes are photographed (with an Hamamatsu camera (Sunayama-cho, Shizuoka, Japan)) and pollen tube length measured with the NIS-element software from Nikon (Nikon Instruments, Melville, NY USA). Pollen tube density and length were also determined through semi *in vivo* analyses. Following hand pollination with wild-type or *larp6c* homozygous mutant pollens, the pistils were excised at the shoulder and incubated for 16 h on a plate containing pollen tube solid growth media. Pistil explants and pollen tubes were photographed, images analyzed with the plugin Bio-format (Open microscopy environment) from the Image J software (Rasband, W.S. ImageJ, NIH, Bethesda USA) (Abràmoff et al. 2004) to determine pollen tube length and density per pistil.

**Ovule attraction assays**

*larp6c-3* or *larp6c-4* pistils were emasculated at stage 12C and left to mature for 2 days. Limited pollination was performed on day 3 using LAT52-GUS homozygous pollen grains in *qrt-/-* background alonside *ms*1-/- pistils as control. Eighteen hours after pollination, pollinated pistils were collected and gently dissected with 25G needle to expose fertilized ovules. Dissected pistils were stained for GUS activity in GUS staining buffer (50 mM PBS, pH7, 0.2% Triton-X100, 10 mM potassium ferrocyanide and 1 mM X-Glu). Pistils were vacuum-infiltrated for 10 min and stained o/n at 37°C. For microscopy analyses, stained pistils were cleared using ethanol series (70%-30%) and mounted on 30% glycerol and analysed with Nikon TE2000 equipped with DIC modulation contrast and NIS-element software. Images were processed with ImageJ.

### SUPPLEMENTAL FIGURE LEGENDS

**Supplemental Figure 1: *LARP6C* mRNA is specifically accumulated in pollen**. (**A**), Expression profiles of the *LARP6A (AT5g46250)*, *6B* (*AT2g43970*) and *6C* (*AT3g19090*) mRNAs across Arabidopsis development. Data were retrieved from the TraVa database of gene expression profiles monitored by RNA-seq (http://travadb.org/) (Klepikova et al. 2015) (upper graph) and from the Arabidopsis eFP Browser from the Bio-Analytic Resources for plant biology (http://bar.utoronto.ca/) (Winter et al. 2007) and represented as histograms. The normalized expression values are arbitrary units. (**B**) RT-PCR analysis of steady-state *LARP6C* mRNA levels in various tissues from wild-type plants. Specificity of the PCR reactions was verified by omitting the reverse transcription step (no RT, lanes 1-4; 9-11). RNAs from *larp6c-3* pollen grains were utilized as negative controls (lanes 1 and 5). RNAs extracted from: seedling aerial parts (lanes 2, 6), whole flower (lanes 3, 7), mature pollen grains (lanes 4, 8), seedling roots (lanes 9, 12), stem (lanes 10, 13) and cauline leaves (lanes 11, 14) were tested. The *ACTIN8* transcript was used as positive control. Supplemental Figure 1 supports Figure 1A.

**Supplemental Figure 2: *larp6c* loss-of-function mutants do not display pollen tube germination nor growth defect. (A)**, *larp6c* loss-of-function induces a slight pollen maturation deficiency. Mature Pollen Grain (MPG) from wild type (WT) or *larp6c-3* homozygous in quartet (*qrt*-/-) mutant background were collected and observed following DAPI staining. The number of bicellular pollens was scored over the total number of observed MPGs. The results are presented as histograms and correspond to mean values of three replicates ± SD and « n » is a total number of pollen grains examined. On the right handside are representative fluorescence micrographs of wild-type and *larp6c-3* pollen grains stained with DAPI. Red arrows point to mature pollen grains in which the generative cell remained undivided and appeared as binucleate. **(B)**, Mean germination rates (± SD) of pollen tubes emerged per pistil in a semi *in vivo* assay. The total number of observed pistils is reported as "n" from three biological replicates. Transgenic LAT52-GFP pollen grains are used as control. **(C)**, Pollen tube length derived from the semi *in vivo* assay. No significant difference was observed between wild-type and mutant pollen tubes (Student *t* test p > 0.05) calculated from pooled replicates. **(D)**, *In vitro* pollen tube length measurements. The total number of pollen tubes measured is reported on the histogram as "n". **(E)**, Semi *in vivo* pollen tube guidance competition assays. On the left, a cartoon representation of pollen tube guidance assay. Unfertilized wild-type ovules were arranged around an *ms1* pistil pollinated both with mutant and LAT52-GFP pollen. A limited pollination was performed with pollen from two different genotypes (as indicated below the graph) to assess the competency of *larp6c-3* and *larp6c-4* mutant pollen tubes. Ovules were scored as: GFP expressing (targeted by LAT52-GFP pollen), non-GFP expressing (targeted by mutant pollen) or non targeted. The results are presented as whisker boxplots. Supplemental Figure 2 supports Figure 1D and E.

**Supplemental Figure 3: Construction of transgenic lines stably expressing tagged versions of LARP6C and subcellular distribution of LARP6C-tRFP.** (**A**) Schematic representation of the transgenes utilized to stably express tagged versions of the LARP6C protein in mature pollen. The drawing and color codes are as in Figure 1B. (**B**) and (**C**), western blot analyses of steady-state levels of tagged versions of the LARP6C protein expressed into the *larp6c-3*, *larp6c-4* or wild-type backgrounds. Total proteins were extracted from whole flowers, and blots probed with anti-LARP6C antibodies. (**B**), total extracts from wild-type (lanes 1) *larp6c-3* (lanes 2) or *larp6c-4* (lane 3) flowers are shown as control. Steady-state levels of the tRFP- or YFP- tagged LARP6C proteins from whole flowers (lanes 4-9). Lines 121 and 1923 (lanes 4 and 8) express fluorescently labelled LARP6C into a wild-type background, lines 134, 136 and 1930 (lanes 5, 6 and 9) in a *larp6c-3* background and line 1885 (lane 7) in a *larp6c-4* background. Confocal analyses presented on Figure 2 were conducted with pollens from: lines 1885 (UNM), 1930 (BCP and TCP), 1923 (MPG) and 136 (pollen tubes). Line 134 was used to co-express the various markers presented in Figure 3 Lines 134 and 1930 were used as complemented lines for pollen experiments reported in Figures 1D and E and Supplemental Figure 2. Black arrows point to endogeneous LARP6C, red arrows to LARP6C-tRFP, green arrows to YFP-LARP6C and black asterisk to aspecific signal. (**C**), total extracts from wild-type (lanes 1) and *larp6c-3* (lanes 2) flowers are shown as control. Steady-state levels of the LARP6C-FlagHA (6C-FH) fusion from six distinct transformants (lanes 3 to 8). The 6C-FH is expressed into *larp6c-3* knockout plants. Mature pollen from line number 267 (lane 5) was used for the RIP-seq experiment. **(D)** Confocal analyses of LARP6C-tRFP distribution in bicellular pollen (BCP), tricellular pollen (TCP) and mature pollen grain (MPG). Line 134 was used to monitor LARP6C-tRFP subcellular distribution. White arrows point to the cytoplasmic connexion (CC), the generative and vegetative cell nuclei (GCN and VCN), the sperm cell nuclei (SCN). Scale bars correspond to 5μM. Supplemental Figure 3 supports Figures 1, 2 and 3.

**Supplemental Figure 4: Ovular attraction competence of *larp6c* lof mutants**. **(A)**, Semi *in vivo* assays using wild-type (WT) (Col 0 ecotype) or *larp6c* (from *larp6c-3* or *larp6c-4* homozygous plants) mutant ovules for wild-type (WT) pollen tube attraction. The histograms report the frequency of targeted/untargeted ovules for each ovule genotype. Error bars are standard deviations calculated from 2 biological replicates with n= 150 ovules (contained into 25 pistil explants) tested for each replicate and each genotype. **(B-F)**, *In vivo* ovule attraction competence assays. Pollen tube targeting events were scored as successful attraction. *ms1*, *larp6c-3* or *larp6c-4* homozygous pistils were pollinated with pollen grains expressing the GUS transgene under the control of the *LAT52* promoter. Ovules showing a positive GUS staining were scored as competent. **(B-E)**, representative micrographs of pollinated *larp6c* mutant ovules (B-D, *larp6c-4*; E, *larp6c-3*). Arrows point to pollen tubes (B and C) and asterisks to pollen tube penetration and burst (C-E). **(F)**, Histogram representation of ovule attraction frequency. The minus GUS (purple) corresponds to frequency of untargeted ovules and the plus GUS (green) to the frequency of targeted ovules. Error bars are standard deviations calculated from 2 biological replicates. The total number of ovules scored in replicate 1 ($n_1$) and 2 ($n_2$) are: *ms1* ($n_1$=802 from 24 pistils; $n_2$= 975 from 29 pistils), *larp6c-3* ($n_1$= 1063 from 31 pistils; $n_2$= 869 from 28 pistils) and *larp6c-4* ($n_1$=860 from 26 pistils, $n_2$= 912 from 29 pistils). Supplemental Figure 4 supports Figure 1C-E.

**Supplemental Figure 5: Validation of the RIP-seq experiment, RIP-Seq filtering workflow and position of A and B boxes on 5'-UTRs of putative LARP6C targets. (A)**, Western blot analysis of the input (lanes 1, 3), unbound (lane 4) and eluate fractions (lanes 2, 5) of the RIP samples. 0.3% (15μL) of the input and unbound and 3% (7.5µL) of the eluate fractions were analysed. The blots were probed with the anti-LARP6C antibody. **(B)**, Comparison of the two biological replicates for the inputs of the RIP-Seq analysis. Correlation of expression levels between replicates of the inputs from wild-type (Col 0) (left panel) or complemented *larp6c-3* ((LARP6C-FlagHA;*larp6c-3*), labelled 6C-FH) (right panel) lines. The X and Y axes correspond to read counts in rpkm respectively for replicates 1 and 2. The expected correlation line for a correlation coefficient of 1 is represented by a red dotted line and the actual correlation line is in black. Correlation coefficients are reported as $R^2$ on each graph. **(C)**, Representation of the workflow utilized to identify the putative mRNA targets of the LARP6C protein in mature pollen grains. The filtering process was conducted with the weighted average values calculated between replicates, and with the list of genes that displayed at least 1 rpkm value in one of the four conditions (inputs, wild type and 6C-FH and eluates, wild type and 6C-FH) (see Table S1). To identify genes Differentially Expressed (DE) between wild type and 6C-FH respectively in input and eluate fractions we used the Cuffdiff suite (Trapnell et al., 2010). The first filtering step consisted in the elimination of all DE genes in the input. To increase stringency, we also filtered out in this first step, genes non-DE but having a Fold Change (FC) values over 1.2, calculated as the ratios between 6C-FH and WT inputs. The second filtering step selects DE genes between the eluate fractions that present a FC of 1 or more, calculated as the ratios between 6C-FH and WT eluates. The third filtering step, is based on the Enrichment Ratio (RE) calculated as the ratio between eluate and input values and firstly selected genes with a RE6CFH>=1. Then we calculated the ratio between RE6C-FH and REWT values, which distribution is represented in **D** and retained genes with a ratio equal or over 3. The target gene list is reported in Table S1. **(E)**, Schematic representation of the 5'-UTRs of Box A and/or Box B containing mRNAs identified as putative LARP6C baits. The positions of the A (red boxes) and B (green boxes) motifs are reported. Correspondence between numbers on the left handside and gene accession numbers are reported in Table S1. Supplemental Figure 5 supports Figure 4.

**Supplemental Figure 6: The LARP6C La-module does not bind the A-type RNA boxes. (A)**, Calorimetric analysis of the interaction between the LARP6C La-module (encompassing residues 137-332) and two A-type oligos (A1 and A2) which sequences are reported below the graphs. For each graph, the upper panel corresponds to the raw titration data showing the thermal effect of injecting an RNA oligos solution into a calorimetric cell containing the recombinant LARP6C La-module. The lower panels show the normalized heat for the titrations obtained by integrating the raw data and subtracting the heat of the RNA dilution. **(B)**, EMSA analyses of LARP6C La-module binding to: A1, A2 or C20 oligos. Decreasing concentrations (μM) 88 (lane 1), 29.3 (lane 2), 9.8 (lane 3), 3.3 (lane 4), 1.1 (lane 5), 0.4 (lane 6), 0.12 (lane 7), 0.04 (lane 8) and 0 (lane 9) of the recombinant La-module were mixed with 3 nM of 5'-labelled oligos. F stands for Free RNA and the red arrows show samples blocked into the wells. Experiments were conducted in the absence (-tRNA) or presence (+tRNA) of unlabelled competitor (tRNAmix of *E. coli* MRE 600 at 0.01 mg/mL concentration). Supplemental Figure 6 supports Figure 4.

**Supplemental Figure 7: Reproducibility of the RNA-seq data.** Comparison of the two biological replicates for the RNA-seq analyses. Correlation of expression levels between replicates of the RNA-seq data from wild-type (Col 0) (left panel) or from *larp6c-3* (right panel) lines. The X and Y axes correspond to read counts in rpkm respectively for replicates 1 and 2. The expected correlation line for a correlation coefficient of 1 is represented by a red dotted line and the actual correlation line is in black. Correlation coefficients are reported as $R^2$ on each graph.

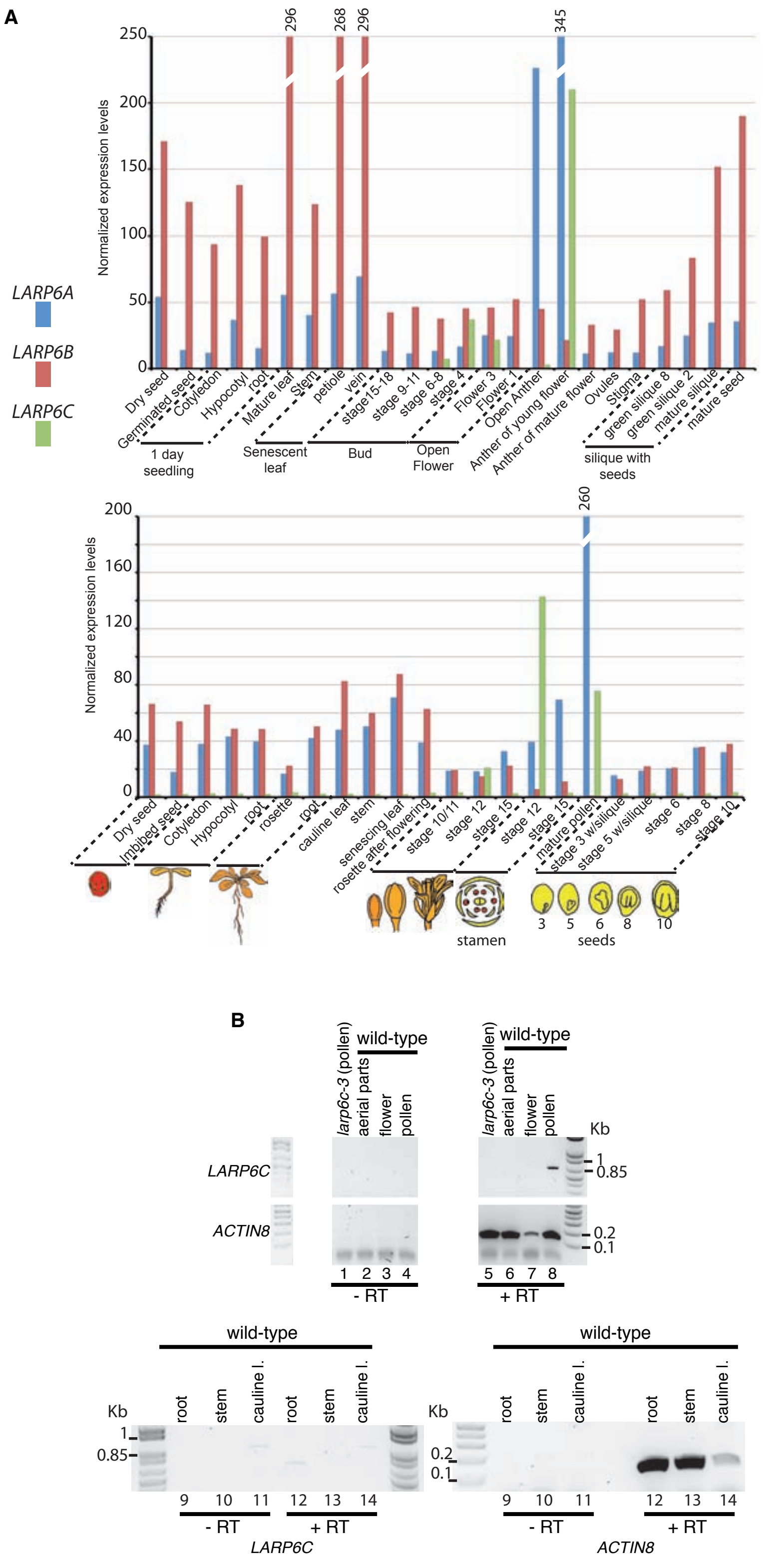


Figure S1
Billey et al.

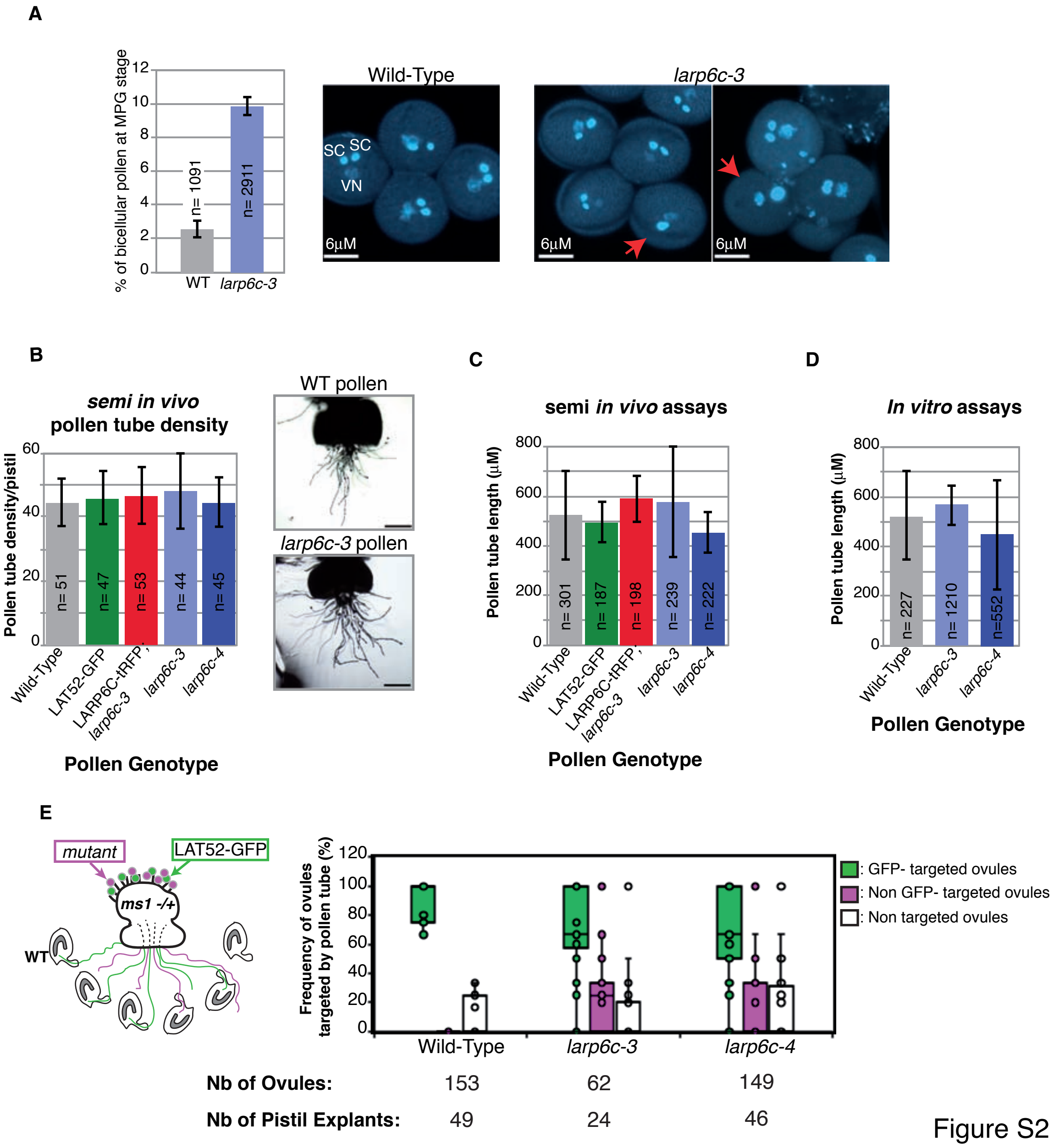


Figure S2
Billey *et al.*

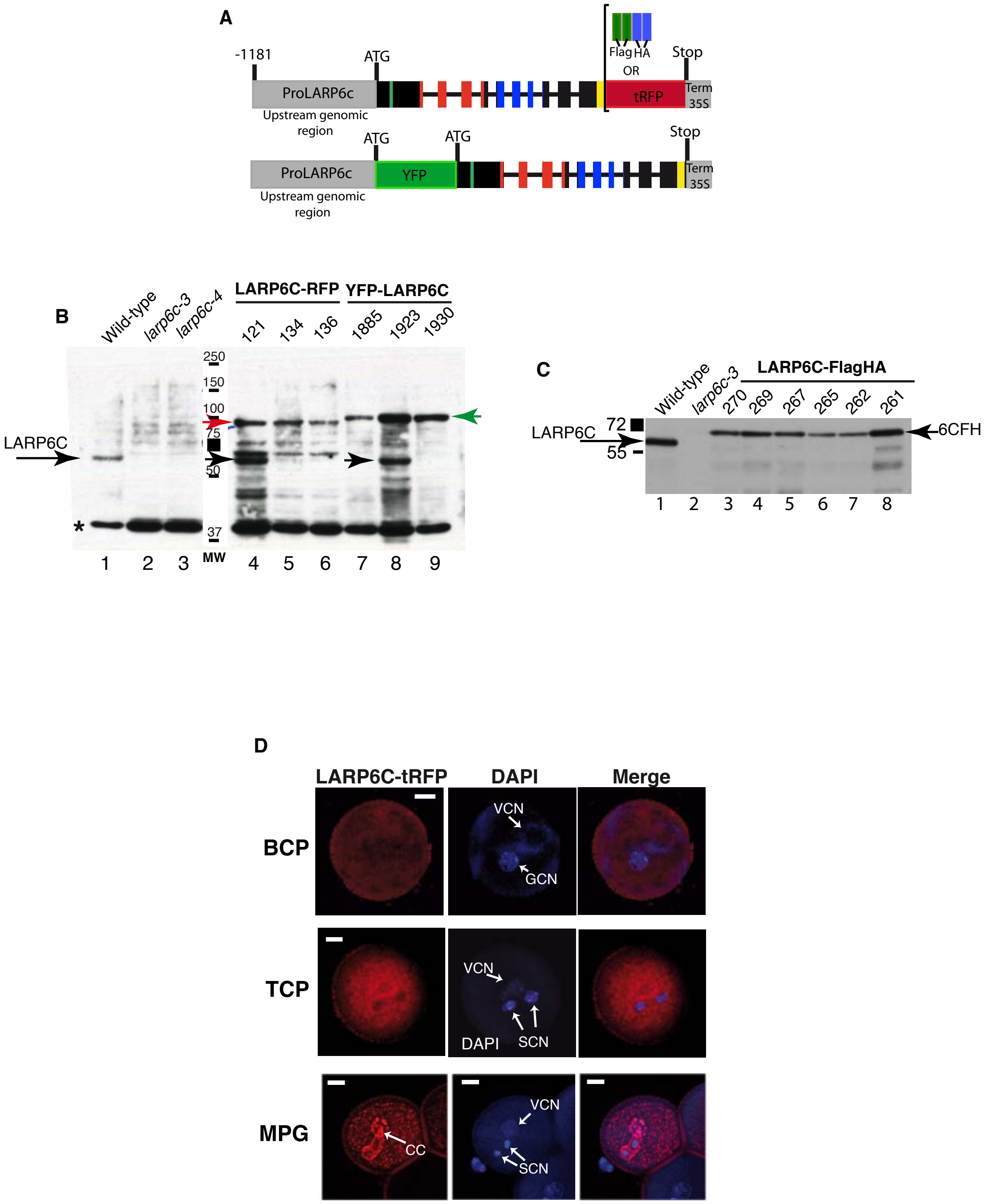


Figure S3
Billey et al.

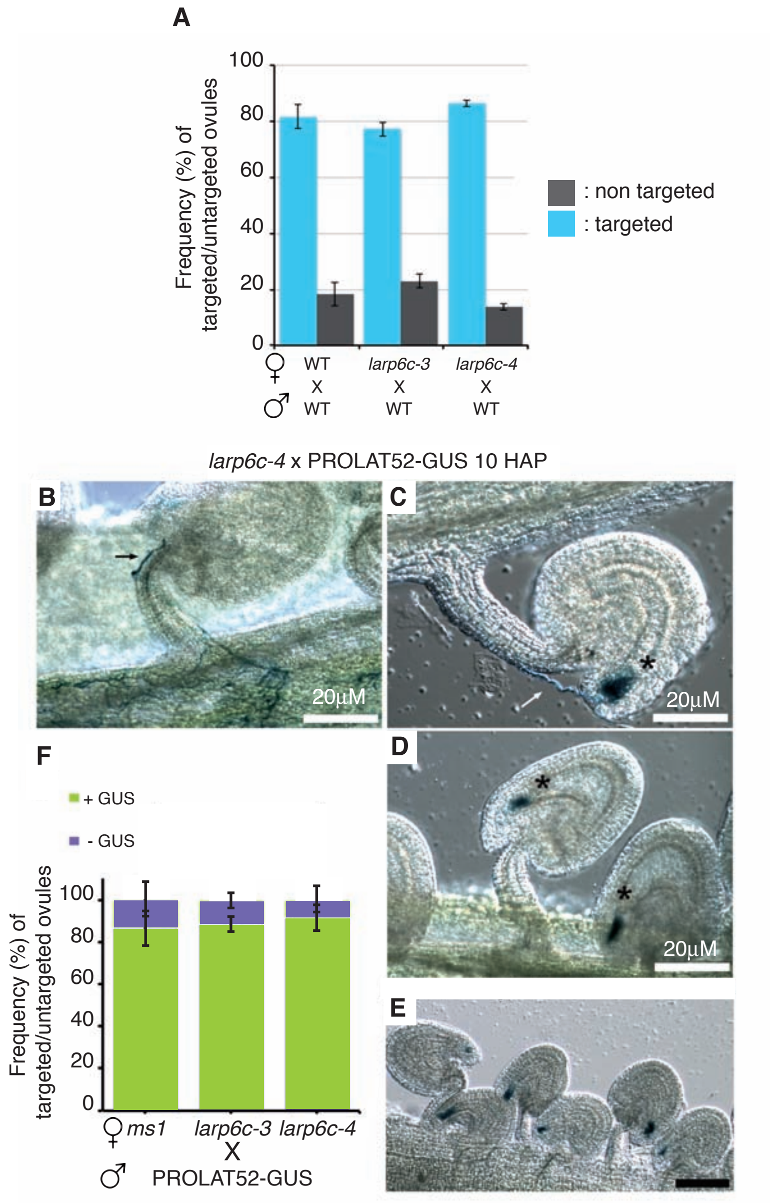


Figure S4
Billey et al

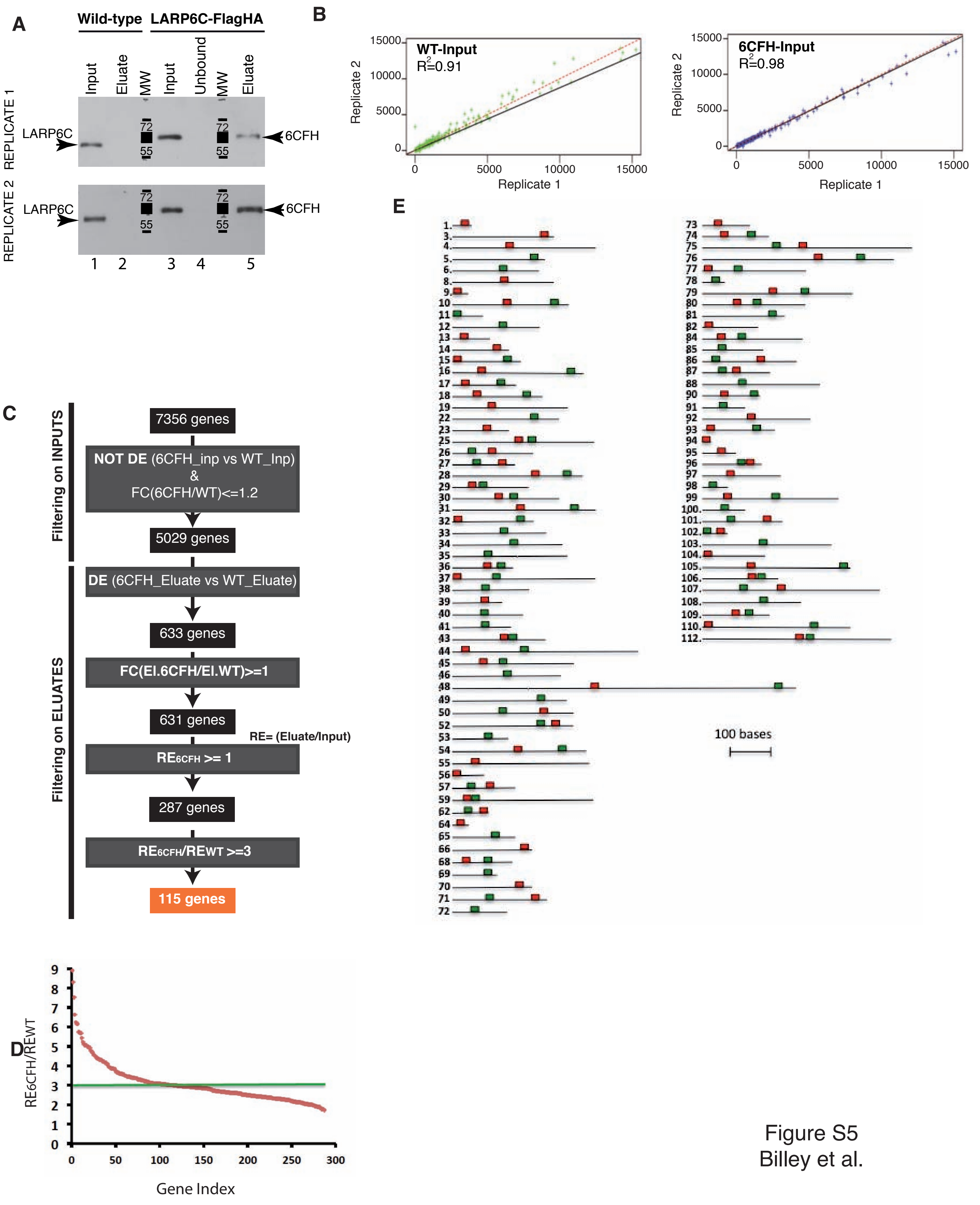


Figure S5
Billey et al.

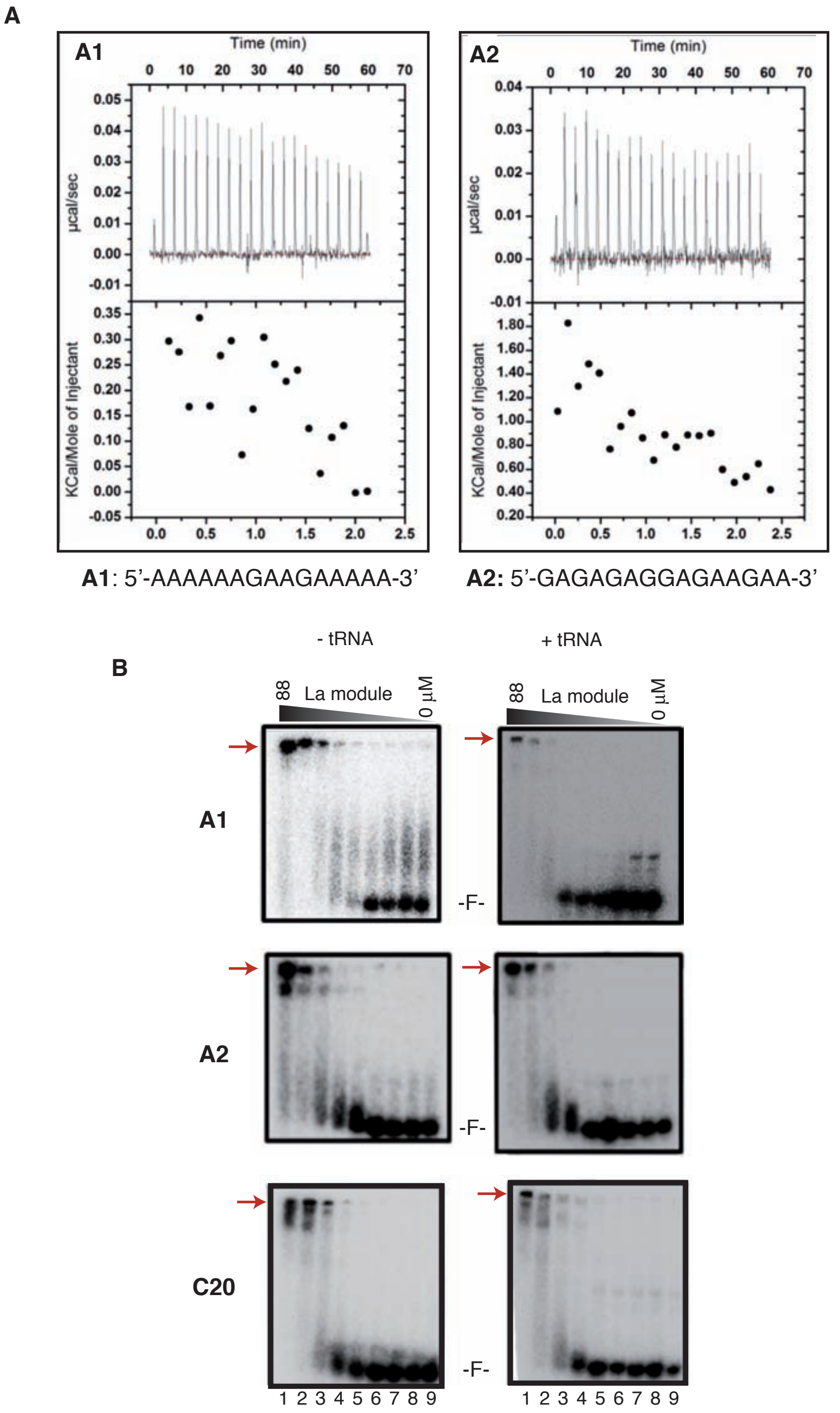


Figure S6
Billey et al.

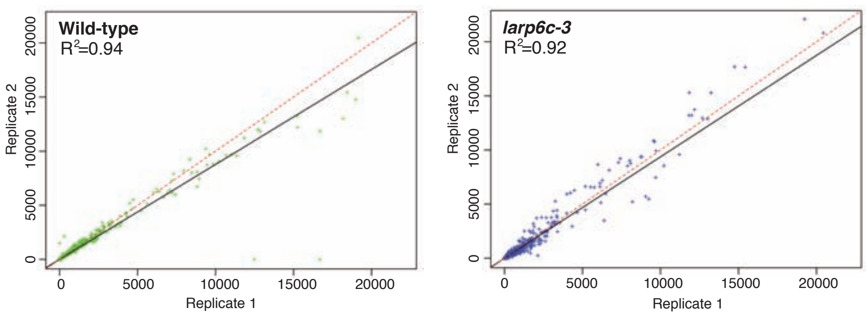


Figure S7
Billey et al.
